# Semaglutide Attenuates Neuroinflammation in Mice

**DOI:** 10.64898/2026.01.10.698763

**Authors:** Dylan M. Rausch, Mette Q. Ludwig, Marie A. Bentsen, Stine N. Hansen, Anna Secher, Dorte Holst, Jaime Moreno, Vivek Das, Kristoffer L. Egerod, Anne-Mette Bjerregaard, Kristoffer Niss, Sarah Bau, Charles Pyke, Kevin Dalgaard, Myrte Merkestein, Franziska Wichern, Charlotte Thim Hansen, Joseph Polex-Wolf, Lotte Bjerre Knudsen, Tune H. Pers

## Abstract

Glucagon-like peptide-1 receptor agonists (GLP-1 RAs) have shown promise in preclinical models of neurodegeneration, with emerging evidence suggesting these effects may be driven by modulation of neuroinflammation. However, the cellular mechanisms underlying GLP-1 RA effects on neuroinflammation remain poorly understood. Here, using a mouse model of lipopolysaccharide-induced neuroinflammation, we investigated how semaglutide coordinates cellular responses to resolve neuroinflammation. We find that semaglutide prevents brain infiltration of neutrophils, excessive cytokine release, and suppresses neuroinflammation-associated transcriptional signatures specifically in microglia, endothelial cells, and a subset of pericytes. Mechanistically, we identify a subset of *Glp1r*-expressing neurons in the dorsal vagal complex that, upon semaglutide treatment, regulate genes involved in anti-inflammatory signaling. Semaglutide-modulated pathways overlap with inflammatory signatures found in human neurodegenerative diseases, including Alzheimer’s disease, suggesting broad relevance for conditions involving neuroinflammation. Together, these findings reveal how GLP-1R signaling orchestrates resolution of neuroinflammation through coordinated multi-cellular programs.

## Introduction

Neuroinflammation is a complex physiological response involving multiple cell types that must be carefully regulated to maintain brain health. While acute neuroinflammation can serve protective functions, dysregulated inflammatory processes are implicated in a host of neurodegenerative diseases, including Alzheimer’s disease (AD)^1,2^. Genome-wide association studies have identified numerous AD risk genes enriched in microglia and immune pathways, and longitudinal studies demonstrate that neuroinflammatory biomarkers predict cognitive decline^3,4^. Interestingly, metabolic disorders such as obesity and type 2 diabetes substantially increase dementia risk, suggesting that metabolic signals may modulate neuroinflammatory processes^5^. However, the cellular and molecular mechanisms through which metabolic pathways regulate neuroinflammation remain poorly understood.

The glucagon-like peptide 1 (GLP-1) signaling pathway has emerged as a key mediator of metabolic-immune communication with neuroprotective properties. GLP-1 receptor agonists (GLP-1 RAs), primarily developed for their metabolic effects^6–8^, demonstrated neuroprotective activity in preclinical models over two decades ago^9^ with subsequent studies revealing anti-inflammatory effects in both central and peripheral tissues^10^. Observational studies in patients with type 2 diabetes suggest that GLP-1 RA treatment may reduce dementia incidence^11–14^. However, recent clinical trials in patients with established AD have not shown cognitive benefits^15^, underscoring the need to understand how, when, and in which cellular contexts GLP-1 receptor (GLP-1R) activation exerts its effects.

Mechanistic work has demonstrated that GLP-1R activation can restore homeostasis in microglia and astrocytes^16–20^, reduce oxidative stress^21–24^, increase blood-brain barrier integrity^19,25,26^, improve vascular health^21,27^, and attenuate peripheral inflammation^21,28–31^. Recent studies have revealed that neuronal *Glp1r* activation in mice is required for the reduction in systemic inflammation through a unique set of neurotransmitter systems^32^ and that brainstem neurons can sense peripheral cytokines and exert control over systemic immune responses^33^, suggesting complex neuro-immune communication circuits.

Despite extensive evidence for GLP-1R’s neuroprotective and anti-inflammatory effects, fundamental gaps remain in understanding how these responses are orchestrated at the cellular level. While microglia are central mediators of neuroinflammation, the requirement for neuronal GLP-1R activation suggests coordination between neurons, glial cells, and vascular components. The specific neuronal populations that express *Glp1r* and mediate anti-inflammatory effects, including brainstem populations implicated in peripheral immune sensing, have not been identified. Similarly, the transcriptional programs engaged in different cell types during inflammation and its resolution, and how these cellular responses relate to disease-relevant signatures in human neurodegeneration, remain uncharacterized. Single-cell genomic approaches now enable systematic investigation of these mechanisms.

Here we investigate how GLP-1R activation coordinates resolution of neuroinflammation using a mouse model of lipopolysaccharide (LPS)-induced neuroinflammation. We created a comprehensive cellular atlas of inflammatory responses and their resolution across vehicle, LPS, and semaglutide treatment using single-nucleus RNA sequencing (snRNA-seq). Our analysis reveals that semaglutide attenuates inflammation-associated transcriptional programs in microglia, endothelial cells, and pericytes while preventing immune cell infiltration. We further identify specific *Glp1r*-expressing neuronal populations in the dorsal vagal complex that activate anti-inflammatory signaling pathways. Importantly, neuroinflammation-associated gene expression signatures are enriched for AD genetic risk loci and recapitulate transcriptional changes in human AD hippocampus, and these signatures are ameliorated by semaglutide treatment. This cell-type-resolved molecular map provides a framework for understanding how GLP-1R signaling coordinates multi-cellular responses to neuroinflammation with implications for therapeutic targeting in neurodegeneration.

## Results

### Semaglutide attenuates LPS-induced neuroinflammation

To investigate how GLP-1 receptor activation modulates inflammatory responses in the brain, we first established a model system allowing detailed temporal analysis of neuroinflammatory processes using LPS. We quantified microglial activation in the hippocampus and the whole brain of 10-week-old C57Bl/6J mice following inflammatory challenge, examining both the early (two days post-challenge; ‘semi-acute’) and resolution phase (11 days post-challenge; ‘sub-chronic’) (n=12 per group; **Fig. 1a**).

**Fig. 1:**
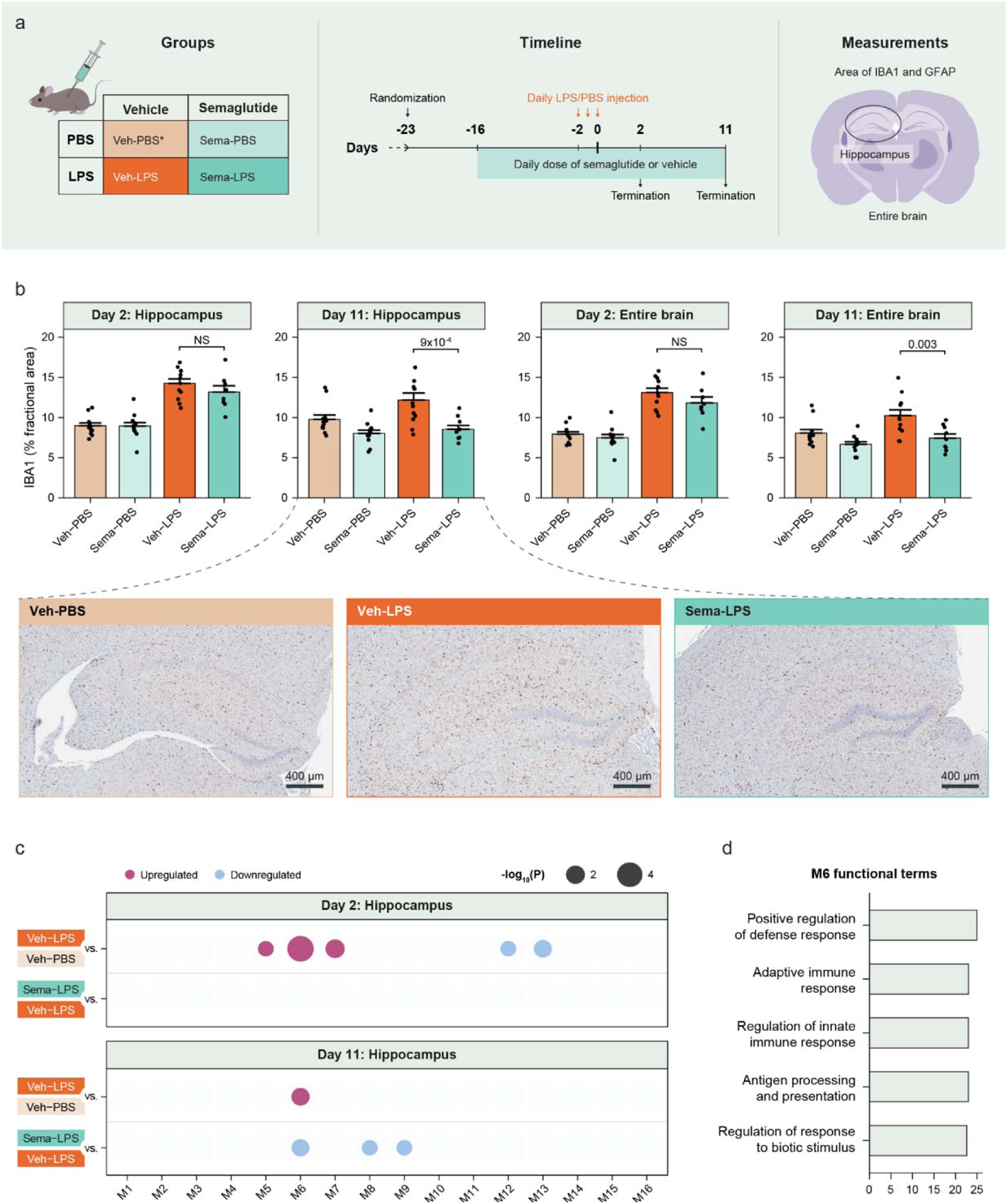
Semaglutide attenuates LPS-induced neuroinflammation at the protein and transcriptional level. **a**, Overview of the *in vivo* study design. Numbers in parentheses denote concentration of LPS. Asterisks indicate the groups represented in the bulk RNA-seq analysis. **b**, Top, relative IBA1 quantified by morphometry (fractional area) in the hippocampus and entire brain on termination 2 and 11 days post-LPS administration, (day 19 and 28 into the study) respectively. Dots represent individual animals and bars, and error bars represent the mean (*n*=8-12) ± SEM. Linear model with BH-adjusted least-squares means two-tailed *t*-test *P.* Bottom, Coronal sections of mouse hippocampi labeled for IBA1using immunohistochemistry 11 days post-LPS administration. **d,** Significantly altered modules of co-expressed genes identified in bulk transcriptomics of the hippocampus from veh-LPS vs. veh-PBS and sema-LPS vs. veh-LPS mice (*n*=6-8 mice/group). Logistic regression with BH-adjusted likelihood ratio test p-value. **e**, The top five most enriched functional terms for co-expressed genes in module M6. BH-adjusted g:Profiler p-value. GFAP, glial fibrillary acidic protein; IBA1, ionized calcium-binding adaptor molecule 1; LPS, lipopolysaccharide; PBS, phosphate-buffered saline; Sema, semaglutide; Veh, vehicle.

Three days of intraperitoneal LPS administration increased microglial activation (measured by IBA1 immunoreactivity) dose-dependently in both hippocampus and whole brain at the semi-acute timepoint. Conversely, only the highest dose of LPS (1.0 mg/kg) induced a sustained increase in area of IBA1 at the sub-chronic time point in the hippocampus (Benjamini-Hochberg (BH)-adjusted *P*=0.011) and entire brain (BH-adjusted *P*=0.006; Supplementary Fig. 1a, 2a). Analysis of astrocyte activation through glial fibrillary acidic protein (GFAP) immunoreactivity revealed coordinated glial responses. The highest inflammatory challenge (1.0 mg/kg LPS) triggered rapid astrocyte activation in both hippocampus (BH-adjusted *P*=0.047) and whole brain (BH-adjusted *P*=4×10^-6^), with sustained activation evident in the whole brain through the sub-chronic phase (BH-adjusted *P*=2×10^-4^; Supplementary Fig. 1b, 2b). This dose and timing provided an optimal window to study both the acute inflammatory response and its resolution.

To assess how GLP-1 receptor activation impacts the resolution of LPS-induced neuroinflammation, we treated mice with semaglutide (30 nmol/kg; n=12 per group) subcutaneously beginning 14 days prior to and throughout LPS administration (**Fig. 1a**). This treatment schedule was designed to minimize the additive appetite-suppressing effects of concurrent LPS and semaglutide administration. Intriguingly, semaglutide’s effects on glial activation showed distinct temporal patterns. While the treatment did not significantly alter early inflammatory responses, it enhanced the resolution of inflammation in the sub-chronic phase, as evidenced by reduced microglial activation in the hippocampus (BH-adjusted *P*=9×10^-4^) and whole brain (BH-adjusted *P*=0.003; **Fig. 1b**). This suggests that GLP-1 receptor activation may primarily promote resolution mechanisms rather than directly suppressing initial inflammatory responses in the CNS.

To define the molecular programs underlying inflammatory resolution, we transcriptionally profiled the hippocampus across all treatment conditions and time points (n=6-8 per group). Using weighted correlation network analysis (WGCNA)^34^, we identified a key gene module that was both upregulated by LPS (BH-adjusted *P*=0.014) and suppressed by semaglutide treatment during the resolution phase (BH-adjusted *P*=0.018, **Fig. 1c**). This module was enriched for genes involved in defense response and adaptive immunity (**Fig. 1d**, Supplementary Table 1), suggesting that GLP-1 receptor activation coordinates a broad program of inflammatory resolution rather than targeting individual inflammatory mediators

### Semaglutide dampens LPS-induced expression changes in glial cells and infiltration of peripheral neutrophils

To map the cell type specific effects of LPS in combination with GLP-1 receptor activation, we performed single-nucleus RNA sequencing (snRNA-seq) of the hippocampus (n=71) spanning the acute (2h, 24h) and sub-chronic (5d, 11d) phases of inflammation (**Fig. 2a**). Using fluorescence-activated nuclei sorting to enrich for non-neuronal cells, we generated a dataset of 109,334 non-neuronal (1,722 unique genes, 3,216 unique transcripts) and 103,607 neuronal cells (3,987 unique genes, 13,696 unique transcripts; **Fig. 2b,c**, Supplementary Fig. 3e). Cell type annotation using reference brain atlases^35–37^ identified all major hippocampal-resident cell populations and, notably, a cluster of infiltrating neutrophils not typically present in healthy brain tissue (**Fig. 2c**, Supplementary Fig. 3a,b). Among the 11 identified neuronal populations (Supplementary Fig. 3c,e), *Glp1r* expression was detected at low levels in two neuronal subtypes – CA3 neurons expressing *Csf2rb2* and caudal ganglionic eminence neurons expressing *Htr3a* (Supplementary Fig. 3d).

**Fig. 2:**
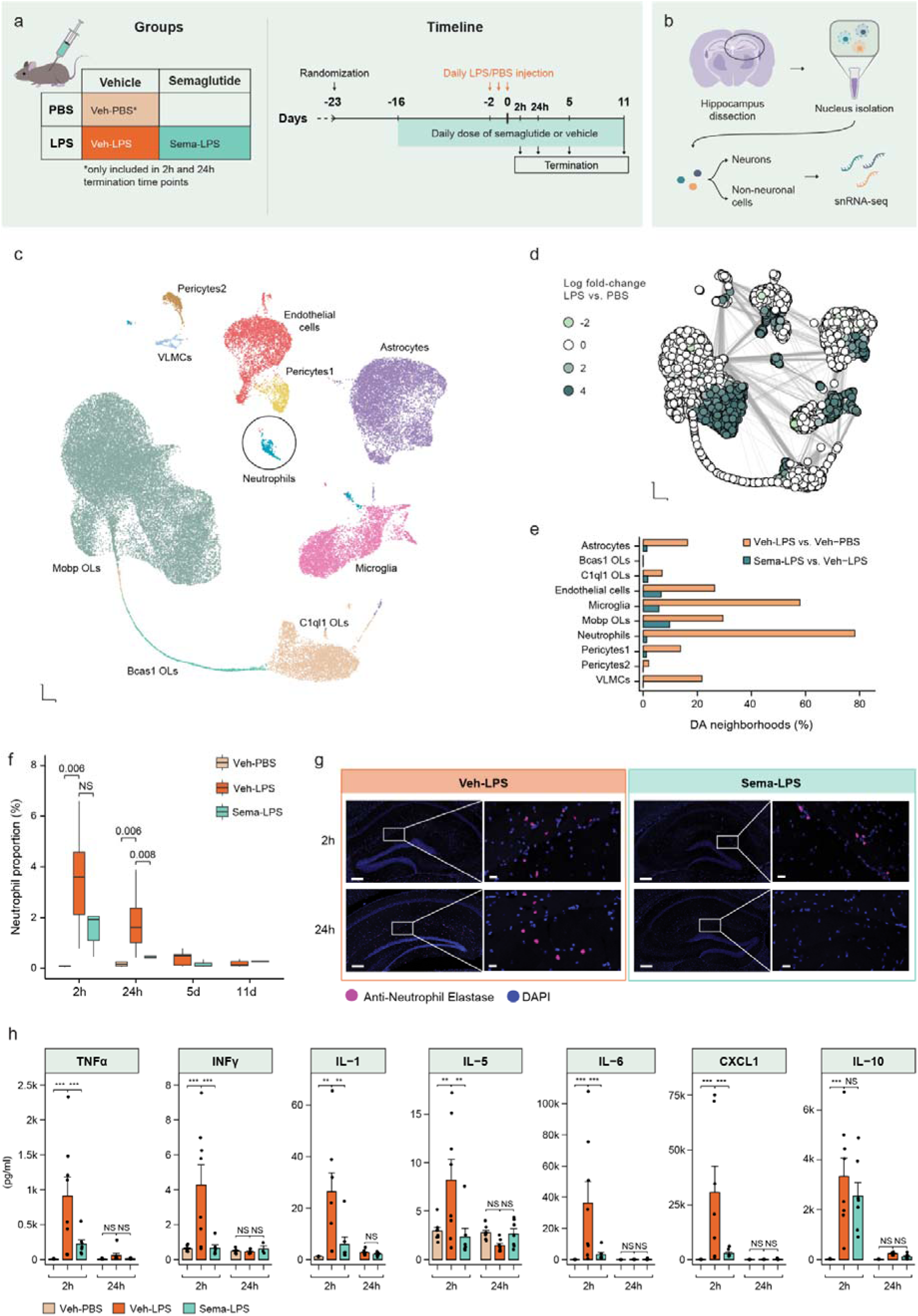
Semaglutide reverses LPS-induced effects in selected hippocampal cell types and peripheral markers of inflammation. **a**, Overview of the in vivo study. *The veh-PBS group was only included in the 2h and 24h termination time points. **b**, Overview of the snRNA-seq protocol of the hippocampus. **c**, UMAP of the 109,334 hippocampal non-neuronal cells colored by the cell type identity. **d**, Neighborhood graph from Milo differential abundance testing of hippocampal non-neuronal cells from veh-LPS vs. veh-PBS mice. Nodes represent neighborhoods of cells. Neighborhoods with significantly more cells from one treatment group are colored by the enrichment of veh-LPS cells relative to veh-PBS. Light and dark green neighborhoods enrich for cells from veh-PBS and veh-LPS mice, respectively. BH-adjusted Milo *P*<0.05. **e**, Percentage of cells assigned to differential abundant neighborhoods (changes induced by LPS and semaglutide treatment in turquoise and red, respectively). **f**, Percentage of neutrophils among all hippocampal non-neuronal cells, shown for each treatment and time point post-administration. Values represent the median, first and third quartiles (Q1 and Q3); whiskers indicate the minimum and maximum values within 1.5 times the interquartile range. BH-adjusted compositional difference *P*. **g**, Representative image pf immunostaining for anti-neutrophil elastase (red) and DAPI (blue) **h**, Terminal plasma levels of TNFα, INFγ, IL-1β, IL-5, IL-6, CXCL1 and IL-10 2h after the final dose of LPS. Dots represent individual animals and bars and error bars represent the mean (*n*=8) + SEM. Unpaired t-test. DA neighborhoods, differential abundant neighborhoods; CXCL1, C-X-C motif chemokine ligand 1; IL-1β, interleukin-1 β; IL-5, interleukin-5; IL-6, interleukin-6; INFγ, interferon gamma; LPS, lipopolysaccharide; PBS, phosphate-buffered saline; Sema, semaglutide; TNFα, tumor necrosis factor α; UMAP, uniform manifold approximation and projections; Veh, vehicle; VLMCs, vascular and leptomeningeal cells.

To identify and prioritize responsive cell populations over time and treatment, we annotated differentially abundant sub-populations of cells using to Milo tool^38^. We found that in general, non-neuronal cells were strongly impacted by LPS during the acute phase (2h and 24h post-LPS; BH-adjusted P<0.05; **Fig. 2d,e**; Supplementary Fig. 3e, 4) with effects diminishing over time. Semaglutide modified the cellular response, introducing Sema-LPS specific cell states in seven out of 10 non-neuronal populations and four neuronal populations (**Fig. 2e**, Supplementary Fig. 3e). Consistent with the Milo analysis and low levels of *Glp1r* expression, semaglutide treatment did not induce gene expression changes in *Csf2rb2* and *Htr3a* neurons (Supplementary Fig 3f). Notably, LPS treatment triggered rapid neutrophil infiltration into the hippocampus that decreased over time, while semaglutide completely prevented this infiltration at 24h post-LPS (**Fig. 2f,g**). Hippocampal immunofluorescence (IF) for anti-neutrophil elastase confirmed that LPS treatment induced hippocampal neutrophil infiltration (P<0.05; n=3) as well as a trend toward an attenuation of infiltration by semaglutide 24h after LPS dosing (Supplementary Fig. 5). These results demonstrate that semaglutide can reduce infiltration of peripheral immune cells while also modulating the transcriptional states of brain-resident cell populations.

Given that central and peripheral immune cells are regulated by a range of cytokines we evaluated the changes in peripheral cytokine levels at two time points following LPS administration. Consistent with previous studies^21,32^, semaglutide significantly reduced multiple pro-inflammatory cytokines 2h post-LPS including tumor necrosis factor alpha (TNFα), interferon gamma (INFγ), interleukin-1 β (IL-1β), interleukin-5 (IL-5), interleukin-6 (IL-6), and C-X-C motif chemokine ligand 1 (CXCL1), while preserving levels of the anti-inflammatory cytokine interleukin-10 (IL-10) (**Fig. 2h**). Cytokine levels correlated with the proportion of neutrophil infiltration, though intragroup variations in cytokine levels did not explain differences in neutrophil abundance (Supplementary Fig. 6). This selective suppression of pro-inflammatory factors while maintaining anti-inflammatory signaling suggests that semaglutide promotes a state of immune tolerance^39^, potentially explaining its effects on both peripheral inflammation and brain immune responses

### Semaglutide activates neurons in the dorsal vagal complex that express inflammation-attenuating neurotransmitters

To determine whether semaglutide’s effects extend beyond the hippocampus, we analyzed the dorsal vagal complex (DVC), a brainstem region that serves as a major access point for semaglutide in the brain^40^. snRNA-seq of DVC tissue from the same experimental groups yielded 141,002 non-neuronal cells and 100,509 neurons (**Fig 3a,b**, Supplementary Fig. 7, Supplementary Table 3). Cell type annotation using a reference atlas^41^ revealed the same non-neuronal populations found in hippocampus plus region-specific ependymal cells and tanycytes (**Fig. 3b**, Supplementary Fig. 7a). Analysis of neuronal populations, which are less discrete in the DVC^41,42^, identified six major classes based on neurotransmitter markers: GABAergic, glutamatergic, catecholaminergic, and cholinergic neurons (Supplementary Fig. 7a,b).

**Fig. 3:**
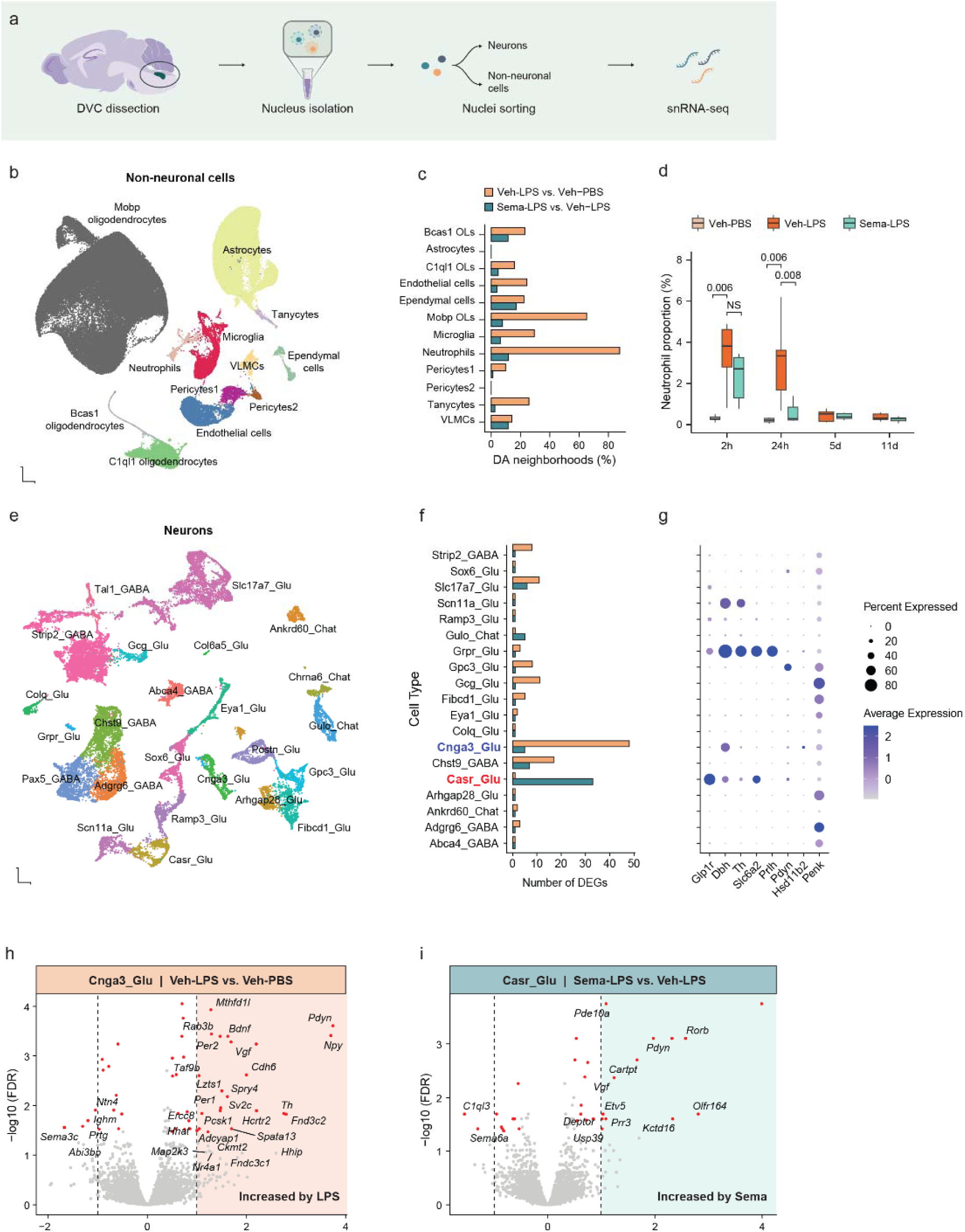
Semaglutide induces transcriptional effects in the DVC indicative of neuron signaling in inflammatory states. **a**, Experimental design. **b**, UMAP of the 141,002 DVC non-neuronal cells colored by the cell type identity. **c**, Percentage of cells of each DVC non-neuronal cell type that are assigned to differential abundant neighborhoods. BH-adjusted Milo *P*<0.05. **d**, Percentage of neutrophils among all DVC non-neuronal cells, shown for each treatment and time point post-administration. Values represent the median, first and third quartiles (Q1 and Q3); whiskers indicate the minimum and maximum values within 1.5 times the interquartile range. **e**, UMAP of the 25 area postrema and nucleus of the solitary tract neuronal populations colored by the cell population identity. **f**, Number of differentially expressed genes identified in each neuronal population in response to LPS (blue) or semaglutide (orange). BH-adjusted DEseq2 *P*<0.05 and absolute log_2_ fold-change>0.5. **g**, Dotplot of the expression of key area postrema and nucleus of the solitary tract neuronal markers. Populations in blue and red are nucleus of the solitary tract- and area postrema-localized respectively. Size indicates percent of cells in a cluster expressing the respective gene and color indicates the expression level. h**-**i, Volcano plot of differentially expressed genes in Cnga3_Glu and Casr_Glu neurons in response to LPS and semaglutide treatment, respectively. DA neighborhoods, differential abundant neighborhoods; DEG, differentially expressed genes; DVC, dorsal vagal complex; LPS, lipopolysaccharide; PBS, phosphate-buffered saline; Sema, semaglutide; UMAP, uniform manifold approximation and projection; Veh, vehicle; VLMCs, vascular and leptomeningeal cells.

Similar to the hippocampus, LPS primarily affected non-neuronal cell populations during the acute phase, with semaglutide modulating these responses (**Fig. 3c**, Supplementary Fig. 8). We again observed neutrophil infiltration specifically in LPS-treated mice, which was attenuated by semaglutide treatment (**Fig. 3d**). These parallel findings in hippocampus and DVC suggest that semaglutide’s ability to prevent immune cell infiltration and modulate neuroinflammation extends across multiple brain regions.

Recent work has shown that GLP-1 RAs require specific neurotransmitter systems to suppress inflammation^32^ and that dopamine β-hydroxylase (*Dbh*)-expressing neurons in the DVC regulate peripheral immune responses to LPS^33^. To investigate potential interactions between these pathways, we extracted and re-clustered neurons from the area postrema and nucleus of the solitary tract (NTS) (**Fig. 3e**, Supplementary Fig. 7b,c). LPS treatment induced substantial transcriptional changes in NTS-resident *Dbh*-expressing neurons (n=47 differentially expressed genes, BH-adjusted *P*<0.05, absolute log_2_ fold-change>0.5), identified here as Cnga3_Glu neurons (**Fig. 3f,g**). Consistent with Jin et al.^33^, LPS upregulated the expression of markers of neuronal activation including *Bdnf*, *Sv2c* and *Vgf* as well as *Pdyn,* a neuropeptide which modulates the inflammatory response (**Fig. 3h**). Importantly, pre-treatment with semaglutide had minimal effects on expression changes in this population (**Fig. 3f**). In contrast, a second population of *Dbh* neurons (Grpr_Glu) that co-express the prolactin releasing hormone (*Prlh*) gene, previously linked to satiety signaling^43^, showed minimal response to LPS (four differentially expressed genes). This selective responsiveness^44^ suggests distinct neural circuits for inflammatory and feeding responses in the DVC.

Area postrema resident *Glp1r*-expressing neurons (Casr_Glu) showed the strongest response to semaglutide treatment, consistent with our previous findings^41^ (**Fig. 3f**). While these neurons were largely unaffected by LPS alone, semaglutide treatment strongly upregulated phosphodiesterase 10a (*Pde10a,* **Fig. 3i**), indicating activation of cAMP-dependent signaling pathways. These neurons co-express *Dbh* and *Slc6a2* (the norepinephrine transporter), and upregulate *Pdyn* upon semaglutide treatment, suggesting increased production of multiple immunomodulatory factors. Together with recent findings, these findings suggest that while LPS and semaglutide act on different neuronal populations, they induce overlapping molecular programs that synergize to modulate immune responses.

### Semaglutide-induced attenuation of neuroinflammation is accompanied by changes in microglia, endothelial cells and pericytes

Given that non-neuronal cells showed the strongest transcriptional responses in both hippocampus and DVC, we next examined whether these responses were consistent across brain regions. We hypothesized that non-neuronal cells may be downstream of the neuronal effects of semaglutide and that the effects of semaglutide and LPS on non-neuronal cells would be similar between the hippocampus and the DVC. Analysis of the top 100 differentially expressed genes for each shared non-neuronal cell type revealed highly similar transcriptional responses between brain regions for both LPS and semaglutide treatment (median Spearman’s rho=0.87 and 0.70, respectively; **Fig. 4a**). Comparing LPS and semaglutide effects within each brain region revealed that semaglutide induced opposing transcriptional changes to LPS (hippocampus: median Spearman’s rho=-0.50; DVC: median Spearman’s rho=-0.49; Supplementary Fig. 9). Together, these results suggest that semaglutide drives a consistent program of inflammatory resolution in non-neuronal cells that counteracts LPS-induced changes across brain regions.

**Fig. 4:**
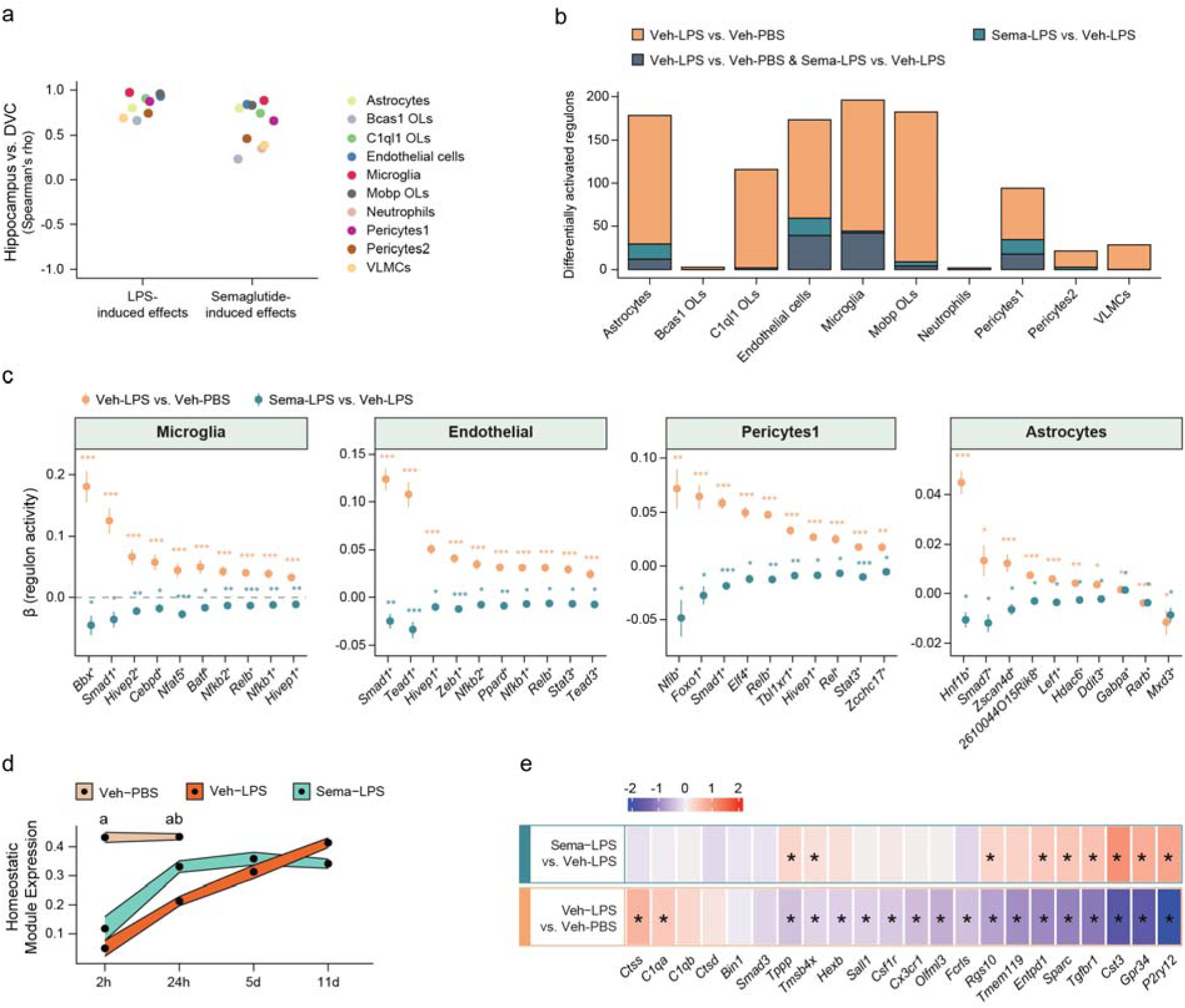
Semaglutide inverses LPS-induced gene expression and gene regulatory networks and promotes a homeostatic state in microglia. **a**, Spearman’s correlation of the log_2_ fold-changes for the top 100 LPS- (veh-LPS vs. veh-PBS) and top 100 semaglutide-induced (sema-LPS vs. veh-LPS) differentially expressed genes between hippocampal and DVC non-neuronal cell types. **b**, Differentially activated regulons comparing cells from veh-LPS vs. veh-PBS mice and sema-LPS vs. veh-LPS mice from the hippocampus. A linear mixed-effects model was constructed, modeling the effect of treatment on regulon activity. BH-adjusted *P* values were computed using a least-squares means two-tailed *t*-test. Regulons with BH-adjusted *P*<0.05 are shown. **c**, Top 10 microglia, pericytes1, endothelial cell, and astrocyte regulons with differential activation in both veh-LPS vs. veh-PBS mice and sema-LPS vs. veh-LPS mice. Regulon transcription factors and linear mixed-effects model β ± SE are shown. Regulons are ordered by the difference in β values between the LPS- and semaglutide-induced effects, and a full list of differentially activated regulons can be found in Supplementary Tables 8-11. **d**, Homeostatic gene expression across time and treatment groups. Linear mixed-effects model and BH-adjusted least-squares means two-tailed t-test *P* (‘a’ indicates *P*<0.05 for veh-LPS and sema-LPS relative to veh-PBS, ‘b’ indicates *P*<0.05 veh-LPS vs. sema-LPS). **e**, Heatmap of selected homeostatic microglia markers as described in Boche and Gordon^47^. Values indicate log_2_ fold-changes of each gene in the comparison. LPS, lipopolysaccharide; PBS, phosphate-buffered saline; Sema, semaglutide; Veh, vehicle; VLMCs, vascular and leptomeningeal cells. *P<0.05, \*\**P*<0.01, \*\*\**P*<0.001.

To identify coordinated transcriptional programs in hippocampal non-neuronal cells, we used the SCENIC tool^45,46^ to map transcription factor networks (henceforth ‘regulons’) across time points. LPS altered regulon activity in eight out of 10 non-neuronal cell types (BH-adjusted *P*<0.05), with the strongest effects in microglia, endothelial cells, astrocytes, and *Mobp*-expressing oligodendrocytes (**Fig. 4b**). Notably, semaglutide treatment inversely regulated all shared regulons in microglia (n=42) and pericytes1 (n=18), and 92% of shared regulons in endothelial cells (36/39) (**Fig. 4b**; Supplementary Tables 4-7). These results demonstrate that semaglutide systematically reverses LPS-induced transcriptional programs in specific non-neuronal cell populations.

Analysis of the inversely regulated regulons revealed cell-type-specific patterns. In microglia, most regulons (30/42) were upregulated by LPS and downregulated by semaglutide (**Fig. 4c**). These regulons were predominantly associated with immune pathways including NF-κB and TNF signaling, while regulons showing the opposite pattern (up with semaglutide, down with LPS) were associated with synaptic membrane and cell adhesion functions (Supplementary Table 8).

A similarly biased pattern was observed in endothelial cells, pericytes1 and astrocytes, but with distinct functional associations (**Fig. 4c**). Pericytes1 regulons primarily involved DNA transcription and cell shape regulation (Supplementary Table 9), while endothelial cells showed a mixed signature overlapping with both microglia (TNF/NF-κB pathways) and pericytes1 (cell shape/transcription; Supplementary Table 8). These patterns suggest cell-type-specific roles in the inflammatory response: immune regulation in microglia, morphological adaptation in pericytes1, and a combination of both in endothelial cells (the astrocyte results can be found in Supplementary Table 11).

Finally, we examined the effect of semaglutide on microglial homeostasis using a consensus transcriptional signature^47^. While this signature remained stable in control animals (**Fig. 4d**), LPS treatment significantly reduced homeostatic gene expression (*P*=1.4×10^-5^). Although semaglutide-treated animals also showed an initial reduction in homeostatic genes (*P*=1.9×10^-4^), they demonstrated significantly faster recovery (BH-adjusted *P*<0.01). Both groups returned to baseline expression by 5d, indicating that semaglutide accelerates restoration of microglial homeostasis. Analysis of individual genes in the homeostatic signature revealed consistent patterns, with *P2ry12*, a receptor controlling microglial motility and inflammatory responses^48^ showing the largest changes (*P*=2×10^-8^; **Fig. 4e**). These results further support our above IBA1-staining-based conclusion that semaglutide promotes a homeostatic immune response.

To examine whether inflammatory resolution involved changes in cell-cell communication, we analyzed ligand-receptor interactions using the CellChat tool^49^. We focused on microglia, endothelial cells, pericytes1, neutrophils and astrocytes as signal-sending cells, while considering all non-neuronal cells as potential receivers, specifically examining interactions involving genes from our LPS-regulated regulons (Supplementary Fig. 10). While neutrophils showed no significant outgoing signals, other cell types displayed distinct interaction patterns. LPS treatment activated multiple microglial signaling programs, including inflammatory regulation (*Batf*), cell morphology and differentiation (*Stat3, Ikzf1, Rad21*), and TNF/NF-κB pathway signaling (*Nfkb1, Rel, Spi1*; Supplementary Table 12). A key interaction involved the *Ptprm-Ptprm* pair, an oxidative stress-sensitive complex controlling barrier function^50^, which featured prominently in TNF and NF-κB-associated regulons. Notably, semaglutide treatment reversed these LPS-induced changes in both gene regulation and cell-cell communication patterns.

### Semaglutide’s attenuating effects on neuroinflammation are supported by human genetics

To examine whether our inflammatory signatures are relevant to human disease, we integrated our data with AD GWAS data. Previous studies have shown that AD-associated genes are predominantly expressed in non-neuronal cells, particularly microglia, peripheral immune cells, astrocytes, and vascular cells^3,51–53^. Initially, to assess whether we could use mouse snRNA-seq data for that same purpose, we assessed whether any of our cell populations enriched for GWAS signals derived from >111,000 clinically diagnosed/proxy AD cases and >677,000 controls^4^. To this end, we first identified the top 1,000 AD-associated risk genes using the MAGMA tool^54^ and then scored each cell across all treatment groups for enrichment of AD-associated GWAS genes using the scDRS tool^55^. Similar to previous studies, we found that both microglia^3,51,52^ and neutrophils^56^ enriched for AD-associated risk genes (BH-adjusted *P*=0.011; **Fig. 5a**). This finding confirms that both cell types express genes linked to AD risk and demonstrates that mouse inflammatory responses involve pathways relevant to human disease^41^.

**Fig. 5:**
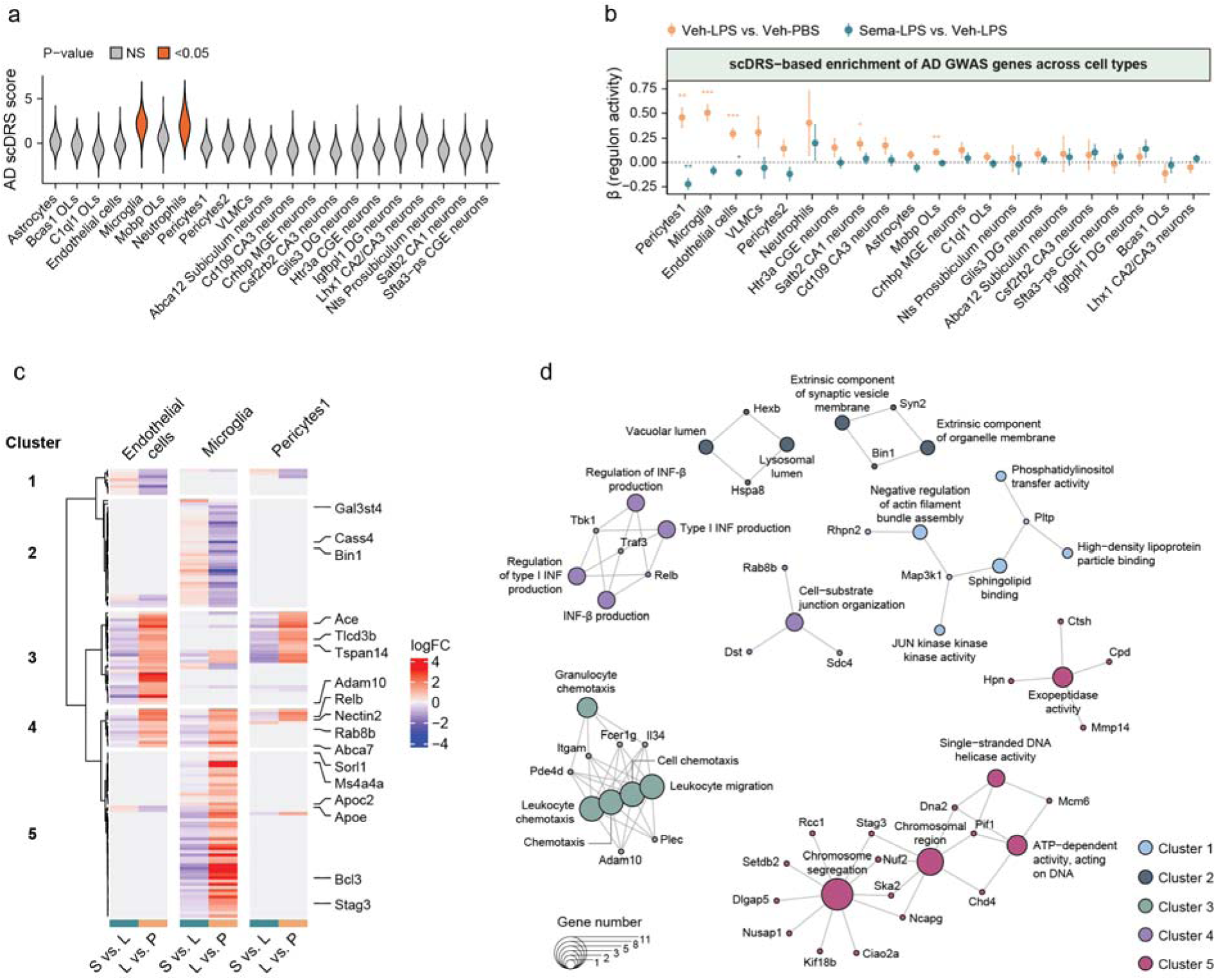
Semaglutide and LPS alter expression of AD GWAS genes. **a**, Violin plot of the scDRS scores for each cell population across all treatment groups. BH-adjusted scDRS *P*. **b**, Comparison of scDRS scores in cells from veh-LPS vs. veh-PBS mice and sema-LPS vs. veh-LPS mice. A linear mixed-effects model was constructed, modeling the effect of treatment on scDRS score for each comparison for each cell population, and BH-adjusted *P* values were computed using a least-squares means two-tailed *t*-test. Linear mixed-effects model β ± SE are shown. Cell populations are ordered by the difference in β values between the LPS- and semaglutide-induced effects. **c**, Expression of 135 regulated AD-associated risk genes in endothelial cells, microglia, and pericytes1 comparing cells from veh-LPS vs. veh-PBS mice and sema-LPS vs. veh-LPS mice (BH-adjusted *P*<0.05). Genes are clustered by the response to treatment and colored by log_2_ fold change. **d**, Gene ontology terms represented as cluster network plot showing specific genes from differentially expressed gene clusters that were present in gene ontology term gene lists (BH-corrected *P*). AD, Alzheimer’s disease; GWAS, genome-wide association study; LPS, lipopolysaccharide; PBS, phosphate-buffered saline; scDRS; single cell disease relevance score; Sema, semaglutide; Veh, vehicle; VLMCs, vascular and leptomeningeal cells. \**P*<0.05, \*\**P*<0.01, \*\*\**P*<0.001.

We next examined whether genes associated with AD risk are regulated during inflammation and its resolution by semaglutide across cell types. We found that LPS treatment increased expression of AD-associated genes in specific cell populations: microglia (BH-adjusted *P*=1.6×10^-8^), pericytes1 (BH-adjusted *P*=1.8×10^-4^), endothelial cells (BH-adjusted *P*=3.1×10^-8^), *Mobp*-expressing oligodendrocytes (BH-adjusted *P*=2.3×10^-4^), and CA1 *Satb2* neurons (BH-adjusted *P*=3.4×10^-3^; **Fig. 5b**). Notably, semaglutide treatment reduced expression of these genes in pericytes1 (BH-adjusted *P*=0.009) and endothelial cells (BH-adjusted *P*=0.032; Supplementary Fig. 11). In microglia, several key inflammatory regulators near the APOE locus^57^ showed opposing regulation by LPS and semaglutide, including *Clptm1*, *Relb*, and *Bcl3* (Supplementary Fig. 11). *Relb* and *Bcl3*, components of the NF-κB pathway^58^, were strongly upregulated by LPS (BH-adjusted *P*=6×10^-13^ and *P*=5×10^-9^) and downregulated by semaglutide (BH-adjusted *P*=0.006 and *P*=0.009). Additional cell type specific changes in AD-associated genes are detailed in Supplementary Table 13. These results reveal substantial overlap between inflammatory pathways and AD-associated genes, suggesting shared molecular mechanisms between acute inflammation and chronic neurological conditions.

To better understand the human relevant pathways are modulated by semaglutide, we analyzed gene expression changes in microglia, endothelial cells, and pericytes, focusing on their overlap with AD-associated genes. We identified 135 differentially expressed genes that overlap with AD GWAS hits, which clustered into five distinct groups based on cell type specific responses (**Fig. 5c**). Microglia-specific responses dominated the pattern, with two clusters (2 and 5) comprising 60% of all regulated genes (87/135). Notably, genes decreased by LPS and restored by semaglutide (Cluster 2) were enriched for lysosomal and vacuolar functions (BH-adjusted P<0.05; **Fig. 5d**), pathways important for both inflammatory responses and cellular homeostasis^59^. Two additional clusters (3 and 4) showed consistent regulation across all three cell types, with genes increased by LPS and reduced by semaglutide. These clusters were enriched for immune cell recruitment (leukocyte chemotaxis, granulocyte migration; BH-adjusted *P*<0.05) and inflammatory signaling (interferon-beta and type 1 interferon production; BH-adjusted *P*<0.01; Fig. 5d), consistent with semaglutide’s effects on neutrophil infiltration and inflammatory resolution. These findings reveal how semaglutide modulates key human disease relevant inflammatory pathways across multiple cell types.

### Semaglutide modulates neuroinflammatory pathways relevant to human disease

Finally, to determine whether semaglutide modulates inflammatory pathways relevant to human disease, we integrated our mouse data with published AD brain transcriptomics (**Fig. 6a**). Analysis of 364 postmortem hippocampi previously identified three molecular subgroups (A, B and C), with Class C characterized by neuroinflammation, increased Aβ load, and APOE4 genotype^60^. In non-neuronal cells, particularly microglia, pericytes1, and endothelial cells, LPS induced transcriptional changes that paralleled those seen in Class C AD patients (**Fig. 6b**). Importantly, semaglutide treatment attenuated these disease-associated signatures across these cell populations and in C1ql1 oligodendrocyte precursor cells, indicating that semaglutide modulates inflammatory pathways relevant to human disease.

**Fig. 6:**
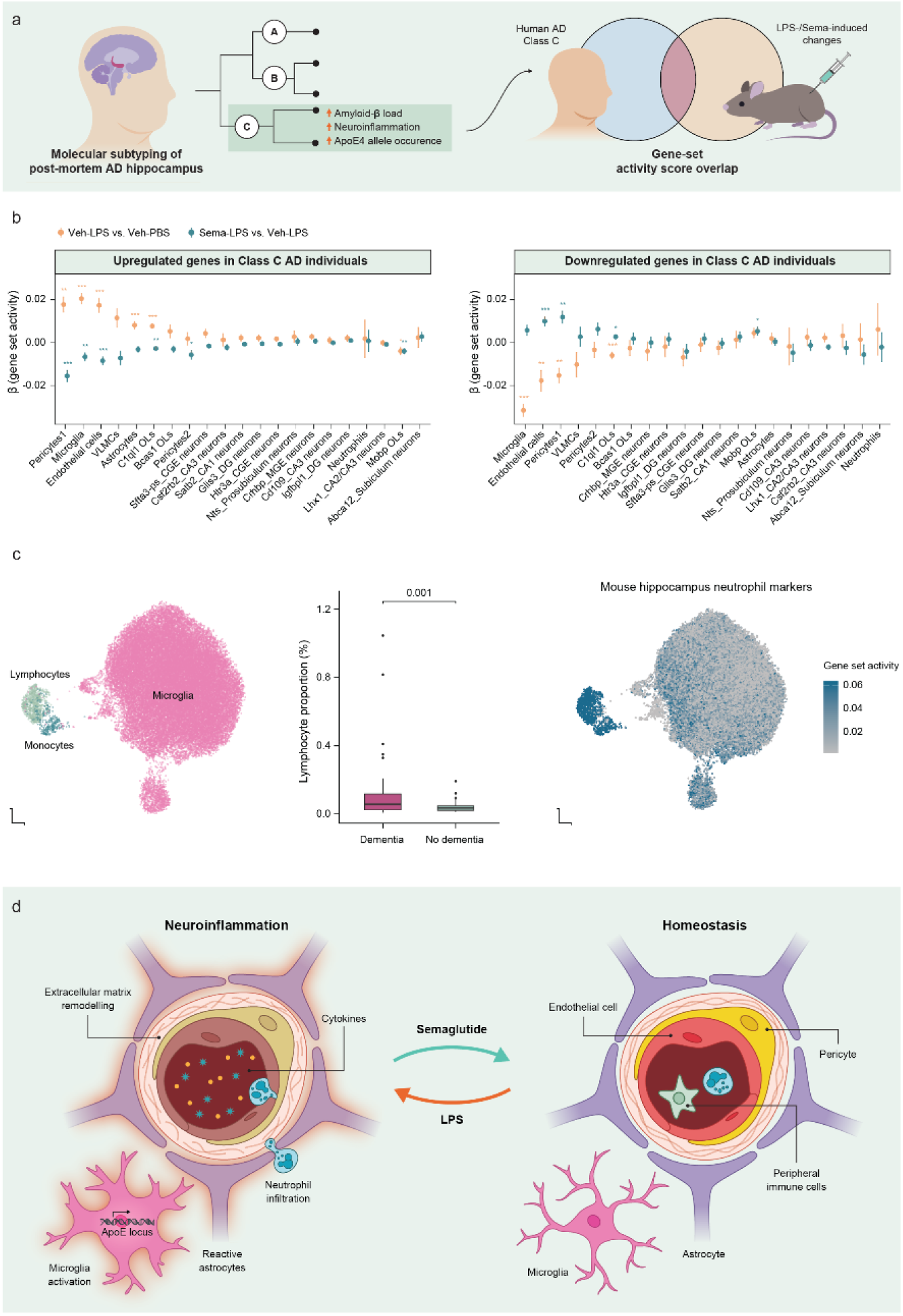
Semaglutide-induced reversal of gene sets identified in human AD and a model of semaglutide-induced attenuation of LPS-induced neuroinflammation. **a**, Genes of molecular Class C was identified based on human postmortem transcriptomics representing Aβ predominant pathology and neuroinflammation^60^ and integrated with snRNA-seq mouse data to compare gene set activity of LPS- and semaglutide-induced changes. **b**, Comparison of activity of Class C AD transcriptional signatures in cells from veh-LPS vs. veh-PBS mice and sema-LPS vs. veh-LPS mice. A linear mixed-effects model was constructed for each comparison for each cell populations, and BH-adjusted *P* values were computed using a least-squares means two-tailed *t*-test. Linear mixed-effects model β ± SE are shown. Cell populations are ordered by the difference in β values between the LPS- and semaglutide-induced effects. **c**, Middle temporal gyrus lymphocyte proportions in human dementia. Left, UMAP of immune cells from the SEA-AD atlas^61^ colored by annotated cell population identity. Middle, Lymphocyte proportion in individuals with and without dementia. Logistic regression with BH-adjusted likelihood-ratio test *P*. Right, UMAP of immune cells colored by mouse neutrophil gene set activity. **d**, Summarized model of how semaglutide restores homeostasis in the setting of LPS-induced neuroinflammation via a multifaceted mechanism, that includes prevention of excessive cytokine release (measured in plasma), reduced infiltration of peripheral immune cells (supported by proportional numbers identified by snRNA-seq), promotion of homeostatic gene expression in microglia, reversal of inflammation-related transcriptional and morphological programs (incl. TNF and NF-κB; WGCNA, IHC, Milo, SCENIC, CellChat) and AD risk genes in immune and cerebrovascular cells (scDRS, gene set activity analysis), change in extracellular matrix gene expression (SCENIC) suggesting extracellular remodeling (grey text). Modest effects of LPS were observed in hippocampal astrocytes (IHC, Milo, SCENIC; grey text). Aβ, amyloid-β; AD, Alzheimer’s disease; IHC, immunohistochemistry; LPS, lipopolysaccharide; NF-κB, Nuclear factor kappa-light-chain-enhancer of activated B cells; PBS, phosphate-buffered saline; scDRS, single cell disease relevance score; Sema, semaglutide; TNF, tumor necrosis factor; Veh, vehicle; VLMCs, vascular and leptomeningeal cells; WGCNA, weighted correlation network analysis. \**P*<0.05, \*\**P*<0.01, \*\*\**P*<0.001.

Given that semaglutide prevented immune cell infiltration in our model, we examined whether similar inflammatory responses occur in human neurodegenerative disease. Analysis of the SEA-AD Brain Cell Atlas, a snRNA-seq dataset from 84 AD patients^61^, revealed infiltrating lymphocytes and monocytes in both middle temporal gyrus and dorsolateral prefrontal cortex. These cells were significantly enriched in tissue from donors with dementia (*P*=0.0012 and *P*=0.0068; **Fig. 6c**, Supplementary Figure 13) and expressed similar markers to the infiltrating cells in our mouse model (**Fig. 6c**). Together, these findings indicate that immune cell infiltration is a conserved feature of brain inflammation, and that semaglutide modulates pathways relevant to human disease.

In sum, our results reveal how GLP-1 receptor activation coordinates resolution of neuroinflammation through multiple mechanisms (**Fig. 6d**). These include suppression of peripheral cytokine release, prevention of immune cell infiltration, and reversal of inflammatory gene programs, particularly TNF and NF-κB pathways that are dysregulated in human disease. At the cellular level, semaglutide promotes restoration of microglial homeostasis and activates specific Glp1r-expressing neurons in the DVC. These neurons express genes involved in anti-inflammatory signaling previously shown to be required for GLP-1 RA’s effects on inflammation. Together, these findings demonstrate how GLP-1 receptor activation coordinates the resolution of neuroinflammation through pathways relevant to human disease.

## Discussion

GLP-1 RAs demonstrate broad anti-inflammatory properties^16–19,21,28–31^, yet the mechanisms coordinating their effects in the brain remain poorly understood. Using immunohistochemistry and snRNA-seq, we demonstrate that semaglutide prevents immune cell infiltration and modulates inflammatory programs across multiple brain cell types. We identify specific *Glp1r*-expressing neurons in the DVC that may coordinate these effects. The inflammatory pathways modulated by semaglutide overlap significantly with those dysregulated in human neurological disease, particularly AD, suggesting broader implications of semaglutide for conditions involving neuroinflammation.

Understanding how GLP-1 receptor activation resolves central and peripheral inflammation has broad implications. Clinical studies demonstrate that GLP-1 RAs can improve brain health outcomes^62,63^ and reduce risk of cognitive decline^11–14^. Although recent trials in established AD did not demonstrate cognitive benefits, semaglutide treatment reduced cerebrospinal fluid levels of YKL-40, several tau species, and neurogranin, biomarkers of neuroinflammation, neuronal injury, and synaptic degeneration, respectively^15,64^. Human genetic studies further support a role for GLP-1 signaling in neuroprotection, with variants affecting GLP-1 receptor function linked to neurological outcomes^65^. The semaglutide-induced attenuation of multicellular inflammatory responses may thus be most effective before irreversible neuronal loss occurs, potentially explaining why biomarker improvements were observed without cognitive benefits in established AD.

Our data suggest several mechanisms through which *Glp1r* activation may reduce neuroinflammation. A key pathway, consistent with findings in other inflammatory conditions^24^, involves modulation of NF-κB and TNF signaling. Specifically, semaglutide suppressed LPS-induced expression of key inflammatory regulators in microglia, including *Relb* and *Bcl3*. The NF-κB pathway, which includes these genes, is central to inflammatory responses and oxidative stress^58^. Here we demonstrate that semaglutide suppresses NF-κB and TNF signaling in both microglia and endothelial cells during inflammatory challenge.

TNF is a potent cytokine released primarily by myeloid cells and elevated TNF expression by microglia is a key mediator of monocyte and neutrophil infiltration in response to LPS^56^. In the current study, semaglutide suppressed both TNF pathway gene expression and plasma TNFα levels, consistent with previous findings in acute inflammation models^21,32^ and clinical studies assessing GLP-1RAs^66,67^. This suppression of TNF signaling likely contributes to the reduced immune cell infiltration and rapid restoration of homeostasis we observed. However, the precise mechanism by which *Glp1r* activation controls TNF and other inflammatory cytokines remains to be determined.

While GLP-1 RAs can act directly on immune cells^16–18^, recent work has revealed that their anti-inflammatory effects require neuronal *Glp1r* activation and downstream adrenergic and opioid signaling^32^. Consistent with this neuronal mechanism, we find that semaglutide induces marked transcriptional changes in *Dbh*-positive, norepinephrine-releasing AP *Glp1r* neurons, among which include *Pdyn*, a precursor for multiple opioid peptides. The requirement for adrenergic receptor signaling points to multiple potential mechanisms through which AP *Glp1r* neurons might contribute to inflammatory responses. Here we demonstrate that semaglutide directly activates neurons that produce endogenous adrenergic and opioid receptor ligands, which could act through *1)* local circuits within the DVC and other brain regions, *2)* efferent nerve fibers innervating immune tissues, *3)* systemic release into circulation, *4)* direct effects on non-neuronal cells, or *5)* a combination of these pathways. The precise mechanism by which *Glp1r* activation engages these neuro-immune circuits remains to be determined.

The prevention of neutrophil infiltration by semaglutide parallels its effects in the lung during polymicrobial sepsis^32^. Several mechanisms might explain this effect. First, semaglutide induction of noradrenergic signaling could regulate vascular function. While we did not detect *Glp1r* expression in hippocampal or DVC vessels (unlike some peripheral tissues^68^), *Glp1r* noradrenergic neurons could modulate vascular tone through adrenergic signaling, potentially affecting neutrophil trafficking and tissue perfusion. Furthermore, our findings of direct transcriptional effects on endothelial cells and pericytes are consistent with a role for GLP-1 RAs in modulating the neurovascular unit itself, potentially contributing to blood-brain barrier integrity and regulating immune cell trafficking independent of direct vascular Glp1r expression. This vascular regulation may be particularly relevant for conditions where vascular dysfunction contributes to pathology, such as diabetic complications and AD^69^. Alternatively, semaglutide might act through efferent nerves to reduce neutrophil mobilization into circulation. Supporting this idea, we found that semaglutide promotes a broader anti-inflammatory environment, reducing pro-inflammatory cytokines while maintaining anti-inflammatory signals like IL-10. This orchestrated response restores immune homeostasis, evident in both microglial morphology and transcriptional state.

The relationship between metabolic disease and neurodegeneration provides important context for these findings. Type 2 diabetes and insulin resistance are established risk factors for AD^70^, with both conditions sharing key pathological features including vascular dysfunction and chronic inflammation^71^. Our observation that semaglutide modulates inflammatory responses across multiple cell types - particularly in microglia, endothelial cells, and pericytes – while maintaining vascular integrity may help explain the emerging clinical benefits of GLP-1 RAs in both metabolic and neurodegenerative conditions^72^.

Despite its strengths, our study has several limitations. First, the mice used in our model were relatively young and hence may have a natural recovery process following LPS that masks additional biological pathways overlapping with AD such as tau pathology. Additionally, semaglutide is known to reduce food intake, and indeed mice treated with semaglutide displayed decreased food intake and body weight loss; however, these effects were close to control before LPS administration and are thus unlikely to explain the effects of semaglutide on peripheral and central inflammation (Supplementary Fig. 10).

In conclusion, our data reveal how *Glp1r* activation orchestrates resolution of neuroinflammation through coordinated effects on multiple cell types. By engaging both neuronal circuits and peripheral immune responses, semaglutide modulates inflammatory pathways that are dysregulated in various neurological conditions. Understanding how GLP-1RAs coordinate these complex cellular responses may provide new strategies for treating neuroinflammatory diseases.

## Methods

### In vivo studies

The care and use of mice in these studies were conducted according to national regulations in Denmark and with experimental licenses granted by the Danish Ministry of Food, Fisheries and Agriculture and the Novo Nordisk Animal Welfare Body. Mice were housed under 12:12 light-dark cycle in humidity- and temperature-controlled rooms with free access to standard chow (catalog 1324, Altromin, Brogaarden, Denmark) and water. Cages were enriched with bedding and nesting material, gnawing sticks, and shelter. A total of 240 male 6-8-weeks old C57BL/6J mice (Janvier, France) were used for the first study (immunohistochemistry (IHC) and bulk RNA-seq) and housed in groups of 3, and 8-10 weeks old at study start (*n* =12/group), and were identified by subcutaneously implanted chips (Pico ID transponder, UNO, OPEND, Denmark). For the second study (snRNA-seq and plasma cytokine), a total of 80 6-8-weeks old male C57BL/6J mice (Charles River, Germany) were singled housed (2 mice/cage separated with divider) upon arrival, and 8-10 weeks old at study start (*n* =7-8/group). All animals were inspected daily by animal caretakers. In both studies, mice were dosed with semaglutide (30 nmol/kg, SC) or vehicle (pH7.4; 0.007 % polysorbate 20; 50 mM phosphate; 70 mM sodium chloride). Mice dosed with semaglutide were uptitrated to the full dose by giving them 3 nmol/kg on the first day of dosing, 10 nmol/kg on the second day of dosing, followed by the full dose of 30nmol/kg daily for the remainder of the study, which is a clinically meaningful dose and within a dynamic dose response ^40^. Mice were dosed with semaglutide or vehicle in the morning between 7-9 am. Two weeks after the onset of dosing with either semaglutide or vehicle, mice were dosed with LPS or PBS for a total of 3 days. On the days of LPS/PBS dosing, animals received their dose of semaglutide or vehicle one hour before the administration of LPS/PBS. In the first study, the dose of LPS was either 0.05 mg/kg, 0.1 mg/kg, 0.5 mg/kg or 1.0 mg/kg daily IP. Mice were euthanized either 2 or 11 days after the last LPS/PBS injection (see Figure 1a). In the second study, the dose of LPS was 1.0 mg/kg daily IP. Mice were euthanized either 2 hours, 24 hours, 5 days or 11 days after the last dose of LPS/PBS (see Figure 2a).

Immediately after termination, terminal plasma was collected, and the whole brain was dissected and divided into left and right hemisphere. For RNAseq samples, the hippocampus was isolated from one hemisphere and placed in liquid nitrogen and stored at -80°C. For IHC procedures, one hemisphere was immersion-fixated in 10% NBF (Neutral Buffered Formalin) for approximately 48h and then transferred to 70% EtOH and stored at 4°C until further use. The hemispheres were paraffin-infiltrated and embedded in blocks. Serial sections representing the rostro-caudal axis of the dorsal hippocampus were cut at 4 μm and collected on Superfrost plus slides. Of the 80 mice used for the single-cell study, three brains were lost, and six hippocampal brain samples had hashing issues and were removed.

### IBA1 and GFAP immunohistochemistry

IBA1 (Abcam, Cat. Ab178845) and GFAP (Dako, Cat. Z0334) IHC was performed using standard procedures. Briefly, after antigen retrieval and blocking of endogenous peroxidase activity, slides were incubated with primary antibody. The primary antibody was detected using a linker secondary antibody followed by amplification using a polymeric HRP-linker antibody conjugate. Next, the primary antibody was visualized with DAB as chromogen. Finally, sections were counterstained in hematoxylin and cover-slipped.

IHC-positive staining was quantified by image analysis using the Visiopharm software (Visiopharm, Denmark). Visiopharm imaging analysis protocols were designed to analyze the virtual slides in two steps:

1. Crude detection of tissue at low magnification (1 x objective) and delineation of Region of Interest (ROI) by artificial intelligence deep learning image analysis to detect hippocampus.
2. Detection of IHC-positive staining and tissue at high magnification (10 x objective) inside the ROI. The quantitative estimates of IHC-positive staining were calculated as an area fraction (AF) in the following way:

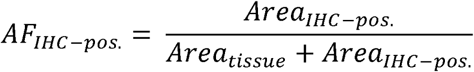

### Anti-neutrophil elastase immunofluorescence staining

Formalin-fixed paraffin-embedded (FFPE) sections of 4.5 µm thickness were obtained and mounted on Superfrost plus slides (Epredia, Cat. J7800AMNZ). Immunostaining for anti-neutrophil elastase (Abcam, Cat. Ab310335) was done using an automated Leica Bond Rx autostainer (Leica Biosystems). The primary antibody was diluted in primary antibody diluent (Triolab, Cat. AR9352) to achieve a final concentration of 1 µg/ml after 30 minutes of heat-induced epitope retrieval using epitope retrieval 2 (Triolab, Cat. AR9640) at 95°C, followed by a 5-minute hydrogen peroxidase blocking step (Advanced Cell Diagnostics, Cat. 322101). The primary antibody was detected using the Brightvision polymeric HRP linker goat anti-rabbit secondary antibody conjugate (Immunologic, Cat. VWRDPVR110HRP) and Opal690 (Akoya Biosciences, Cat. FP1497001KT) diluted in RNAscope Multiplex Tyramide Signal Amplification buffer (Advanced Cell Diagnostics, Cat. 322809). Subsequently, the sections were counterstained with DAPI (Advanced Cell Diagnostics, Cat. 322764) and mounted with Prolong Diamond antifade mounting medium (Invitrogen, Cat. P36970). Fluorescent images were captured using the VS200 digital slide scanner (Olympus) employing a UPLXAPO20X (NA 0.8) air objective with consistent exposure settings. Image processing for publication was conducted using Olympus OlyVIA. The scale bar size is provided in the figure legends. Image analysis for the quantification of hippocampal neutrophils was performed using Halo software (Indica Labs).

### Bulk RNA-seq of the hippocampus

Samples were stored at -80°C until processing. RNA was isolated using the NucleoSpin® kit (MACHEREY-NAGEL). A total of 10 ng-1 μg purified RNA from each sample was used to generate a cDNA library using the NEBNext® UltraTM II Directional RNA Library Prep Kit for Illumina (New England Biolabs). cDNA libraries were then sequenced on a NextSeq 500 using NextSeq 500/550 High Output Kit V2 (Illumina).

### Plasma cytokines

Plasma levels of TNFα, IFNγ, IL-1β, IL-5, IL-6, CXCL1, and IL-10 were analyzed using a mouse multiplex Meso Scale Discovery platform (Meso Scale Diagnostics, Rockville, Maryland) according to the manufacturer’s instructions.

### Single-nucleus RNA-seq of the hippocampus with NeuN depletion

Frozen tissues from the hippocampi were mechanically dissociated in a 2mL glass tissue douncer (Sigma, cat. no.: D8938 with pestels) using 1mL Nuclei EZ Lysis Buffer (Sigma, cat. no.: NUC101-1KT) and 10 firm strokes with glass pestle B. Subsequently, the homogenized samples were incubated on ice for 5 min and transferredthrough a 40µM cell strainer (PluriSelect, cat. no.: 43-10040-40) to a 2mL Protein LoBind tube (Sigma, cat. no.: EP0030108132). The samples were centrifuged (Eppendorf 5810R) for 10 min at 1000*g* with brake set to 1 and resuspended in 250µL Nuclei Buffer (1% BSA (Sigma, cat. no.: SRE0036), 2mM MgCl_2_ (Sigma, cat. no.: M1028), 0.04U/µL Protector RNase inhibitor (Sigma, cat. no.: 3335399001) in PBS, pH 7.4 w/o Ca^2+^, Mg^2+^ (Gibco, cat. no.: D1556)). The samples were further subjected to an Iodixanol gradient purification, where 250µL of 50% Iodixanol/nuclease-free water (Sigma, cat. no.: D1556) was added, and the samples were thoroughly, but gently, mixed by pipetting and layered on top of 500µL 29% Iodixanol followed by centrifugation for 22min at 14,000g with brake set to 1. The isolated nuclei were located at the bottom of the tube and resuspended in 500µL Nuclei Buffer and incubated on ice for 15 min.

Samples were centrifuged for 10 min at 1000g with brake set to 1 and resuspended in 100µL Nuclei Buffer with 0.5µg TotalSeq^TM^-A anti-Nuclear Hashtag (HTO) for multiplexing and 0.5µg NeuN Alexa Flour 488 (Millipore, cat. No.: MAB377X). Samples were then transferred to a 1.5mL Protein LoBind tube (Sigma, cat. No.: EP0030108116) and incubated on ice for 30 min. Samples were again centrifuged for 10 min at 1000g with brake set to 1 after which samples were resuspended in 200µL Nuclei Buffer with 0.4µg 7-AAD (Sigma, cat. No.: 3462) for sorting.

From each sample we aimed to 4,000 7-AAD-positive nuclei (SONY SH800S cell sorter) using a 70µm sorting chip (Sony Biotechnologies, cat. No.: LE-C3207) into a 2mL Protein LoBind tube with 18.8µL RT Reagent B (10X Genomics, Chromium Next GEM Single Cell 3’ Kit v3.1, cat. No.: PN-1000268). Of the 4,000 sorted nuclei, 2,000 were NeuN^low^. Following sorting, the volume was adjusted to 43.1µL with Nuclei Buffer and the final GEM Master Mix reagents were added as per manufacturer’s procedure, which was followed from then on for library preparation with dual indexing.

HTO libraries were prepared by following the procedure from BioLegend and quantified by Qubit and TapeStation (Agilent TapeStation 4200 System) with TapeStation High Sensitivity D1000 DNA (Agilent, cat. no.: 5067-5584 and 5067-5585).

### Single-nucleus RNA-seq of the DVC

Frozen tissues from DVCs were isolated as described for the hippocampal samples and resuspended in 500µL Nuclei Buffer without subsequent Iodixanol gradient purification.

Samples were then centrifugated as described for the hippocampal samples and resuspended in 100µL Nuclei Buffer with 0.5µg TotalSeq^TM^-A anti-Nuclear Hashtag antibody for multiplexing^73^, transferred to a 1.5mL Protein LoBind tube (Sigma, cat. no.: EP0030108116) and incubated on ice for 30 min. Following centrifugation (as above), the sample was resuspended in 200µL Nuclei Buffer with 0.4µg 7-AAD (Sigma, cat. no.: 3462) for sorting.

From each sample, 2,500 7-AAD-positive nuclei were sorted (SONY SH800S cell sorter) with a 70µm sorting chip (Sony Biotechnologies, cat. no.: LE-C3207) into a 2mL Protein LoBind tube with 18.8µL RT Reagent B (10X Genomics, Chromium Next GEM Single Cell 3’ Kit v3.1, cat. no.: PN-1000268). Following sorting, the volume was adjusted to 43.1µL with Nuclei Buffer and the final GEM Master Mix reagents were added as per manufacturer’s procedure, which was followed from then on for library preparation with dual indexing.

HTO libraries were prepared as described for the hippocampal samples.

### Bulk RNA-seq raw data processing

The bulk RNA-seq data was aligned to the mouse genome obtained from the Ensembl database using the Spliced Transcripts Alignment to a Reference (STAR) software.

### Single-nucleus RNA-seq raw data processing

BCL files were demultiplexed into FASTQ files using bcl2fastq v.2.19.01 ^76^. Reads were pseudoaligned using Salmon Alevin v.1.9.02 ^77^ with the flags *--read-geometry 2[1-15] --bc-geometry 1[1-16] --umi-geometry 1[17-26]* set. The RNA library was pseudoaligned to the GENCODE vM23 reference transcriptome, distinguishing between spliced and unspliced transcripts. Alevin-fry v.0.7.03 ^78^ was used to quantify both the RNA and HTO libraries. Quantified libraries were subsequently processed with Seurat ^79^. Empty droplets were detected with the *barcodeRanks* function from the DropletUtils package ^80^. HTOs were normalized using the *NormalizeData* function with the normalization method set to *CLR*. A Gaussian mixture model with two components was fitted to each HTO distribution and a droplet was called to be positive for this HTO if it was predicted to belong to the cluster with the higher mean expression. Droplets positive for multiple HTO were classified as doublets. Inter HTO doublets were found using the *recoverDoublets* function form the scDblFinder package ^81^. Cell negative for HTO library as well as doublets were removed.

### Bulk RNA-seq data analysis

#### WGCNA

WGCNA was run using the R implementation. Raw counts were subjected to variance stabilizing transformation (vst) normalization using DESeq2 and transformed into a similarity matrix using the biweight midcorrelation. From the similarity matrix, a signed network was constructed using a soft-thresholding power of four, maximizing the scale-free topology *R*^2^ fit. Subsequently, the network was subjected to hierarchical clustering based on the average topological overlap measure, and modules of co-expressed genes were identified using the *cutreeDynamic* function with the parameters minimum cluster size set to 30, deep split set to three, and pam stage set to false. Finally, module eigengenes were calculated and modules with a Pearson correlation above 0.75 were merged.

Modules with altered activity following treatment were identified using a logistic regression framework. For each module, a logistic model with the module eigengene as the dependent variable and the treatment group as independent was constructed and compared to the null model using a likelihood-ratio test. *P* values were corrected for multiple testing using the BH method (adjusting for the number of modules).

#### Functional gene set enrichment analysis (WGCNA)

Functional gene set enrichment analyses were carried out using the g:Profiler R implementation. Module genes were ordered by their kME (module membership) values and used as input for the *gost* function with sources set to GO (molecular functions, biological processes, and cellular components), KEGG, and REACTOME terms, ordered query set to true, and correction method set to BH. Terms with more than 500 genes were not considered. Similarly, terms with less than three intersecting genes were excluded.

### Single-nucleus RNA-seq data analysis

#### Initial processing

Downstream analysis was carried out in Seurat. For each batch, cells with outlier UMI counts were identified and removed. Additionally, cells with a mitochondrial RNA content >1% were discarded. Raw counts were normalized with the *SCTransform* function, and principal component analysis (PCA) was performed with the *RunPCA* function using the top 3,000 variable genes as input. Subsequently, clustering was performed with the *FindNeighbors* and *FindClusters* functions using the top 30 PCs as input.

For both the hippocampus and DVC, cells were split into neurons and non-neuronal cells based on the expression of known marker genes. Normalization, PCA, and clustering were rerun as described above for the neuronal and non-neuronal atlas separately. To identify clusters of low-quality cells, for the neuronal atlas, for each batch and cluster, the mean mitochondrial RNA content and the mean expression of *Mobp* (marker of oligodendrocytes; the most abundant non-neuronal cell population) were computed. Clusters with a mean above the 95% quantile in all batches were removed. Additionally, clusters where more than 75% of cells originated from one batch were discarded. For the non-neuronal atlas, for each batch and cluster, the mean mitochondrial RNA content and the mean expression of *Rbfox3* (marker of neurons) were computed, and clusters with a mean above the 95% quantile were likewise discarded.

#### Hippocampal cell population annotation

To group hippocampal cells into cell populations, clustering was run on the final neuronal and non-neuronal atlas with the *FindNeighbors* and *FindClusters* functions. To identify an optimal resolution, clustering was run at 10 different resolutions (0.1-1), and for each resolution, the median silhouette score was computed. For both the neuronal and non-neuronal atlas, a resolution of 0.1 yielded the highest median silhouette score (0.62 and 0.47 for the neuronal and non-neuronal cells, respectively).

To annotate the neuronal and non-neuronal cell populations, labels were transferred from published mouse hippocampal atlases. For the neuronal atlas, an atlas consisting of ∼1.3 million neurons from the isocortex and hippocampal formation was utilized as reference^37^. These cells were profiled with either the 10x Genomics or Smart-seq platform and were clustered into 388 neuronal populations assigned to 42 subclasses. To limit computational costs, only cells profiled with the Smart-seq platform (∼73,000 cells) were used. The *FindTransferAnchor* and *TransferData* functions were applied to transfer the subclass labels from cells originating from the hippocampus.

To annotate the non-neuronal atlas, an atlas consisting of ∼20,000 cells from the hippocampus profiled with the 10x Genomics platform ^35^ was used. The *FindTransferAnchor* and *TransferData* functions were utilized to transfer the labels from cells classified as non-neuronal cells. In the non-neuronal atlas, one cluster comprised cells that expressed genes involved in immune function that were distinct from microglia and did not share markers with a specific non-neuronal cell population. To annotate these cells, an atlas of immune cells from the meninges comprising ∼131,000 cells across seven different studies ^36^ was applied. Based on this label transfer approach, three clusters of non-neuronal cells were split into two different cell populations, respectively: 1) pericytes2 and VLMCs; 2) endothelial cells and pericytes1; and 3) *Bcas1* and *C1ql1* oligodendrocytes were originally assigned to one cluster each. As the median silhouette score of this updated clustering (0.61) was higher than for the original clustering, the three clusters were each split into two clusters in the final atlas.

#### DVC cell population annotation

To group DVC cells into cell populations, clustering was run on the final neuronal and non-neuronal atlas with the *FindNeighbors* and *FindClusters* functions. To annotate the non-neuronal cell populations, clustering was run at a resolution of 0.1 and 1, and labels were transferred from a published AP-centric DVC atlas ^41^.

To annotate the neuronal cell populations, clustering was run at a resolution of 1, and clusters were merged and labeled based on the expression of markers of key neurotransmitters (*Slc32a1*, *Gad2*, *Slc17a6*, *Chat*, *Dbh*, *Th*, and *Tph2*). To extract and subcluster neuronal populations originating from the area postrema and neighboring nucleus of the solitary tract, the DVC neuronal atlas was integrated with the published AP-centric neuronal atlas ^41^ using the *SelectIntegrationFeatures*, *PrepSCTIntegration*, *FindIntegrationAnchors*, and *IntegrateData* functions. Following PCA dimensionality reduction, the integrated atlas was clustered at a resolution of 1 and 2. Cells belonging to resolution 1 clusters, where less than 20% of the published AP-centric neuronal populations mapped to were removed from the DVC neuronal atlas. Likewise, cells belonging to resolution 2 clusters, where less than 10% of the published AP-centric neuronal populations mapped to were removed from the DVC neuronal atlas. This updated AP-centric DVC neuronal atlas was subsequently labeled based on the mapped published AP-centric neuronal populations, and PCA and UMAP dimensionality reduction was rerun.

#### Milo analysis

Differential abundance of cellular neighborhoods across treatment was assessed using the Milo tool v.0.99.6 ^38^. The Milo framework was run separately on cells from veh-PBS and veh-LPS animals and cells from veh-LPS and sema-LPS animals. Initially, PCA was recomputed, and the Seurat object was turned into a Milo object with the *as.SingleCellExperiment* and *Milo* functions. A neighborhood graph was built with the *buildGrpah* function using the top 30 PCs as input, considering the top 50 nearest neighbors. Representative neighborhoods were subsequently identified using the *makeNhoods* function, and the distances between neighborhoods were computed with the *calcNhoodDistance*. Differential abundance analysis was performed by running the *countCells* function to count the number of cells in each neighborhood followed by the *testNhood*s function to test for differential abundance between cells from different treatment groups, taking the interdependence between cells originating from the same animal into account. The *P* values were corrected for multiple testing using the spatial BH method implemented in Milo (adjusting for the number of neighborhoods). Finally, to assess the percentage of differential abundant neighborhoods within each cell type, for each neighborhood, the cell type identity was defined as the most abundant cell type, with the exception that neighborhoods with less than 70% cells labeled as the same cell type being defined as a ‘Mixed’ neighborhood.

#### Analysis of compositional differences

Compositional differences of neutrophils across treatment and time were assessed using the Cacoa tool v.0.3.0 ^82^. The *estimateCellLoadings* function was run using the non-neuronal cells from veh-PBS and veh-LPS animals and non-neuronal cells from veh-LPS and sema-LPS animals as input, respectively. The *P* values for changes in neutrophil loadings were then corrected for multiple testing using the BH method (adjusting for the number of treatment group comparisons and time points).

#### Differential gene expression analysis

For differential gene expression analysis, a pseudo-bulk count matrix was generated by summing the transcript counts for all cells within the same cell type and mouse identity combination. EdgeR v4.4.0 was used to identify DEG. Treatment group and time point post-PBS or LPS treatment were included as variables in the design matrix. P-values were corrected for multiple testing using the BH method.

#### Functional gene set enrichment analysis (DE)

Functional gene set enrichment analysis was carried out using the clusterProfiler (version 4.4.4) and org.Mm.eg.db R packages to further explore the functional role of the clustered differentially expressed AD GWAS genes. GO enrichment was run on all 5 DEG cluster lists together using the *compareCluster* function in clusterProfiler with the “enrichGO” option and a *P* value cutoff□<□0.01 and BH-adjusted *P* value cutoff□<□0.05 against all ontologies. Cnetplot functions within the clusterProfiler package was used to create visualizations of the significant GO enrichments.

#### Microglia Homeostatic Gene Expression

A homeostatic gene expression score was computed using the AddModuleScore() function in Seurat. For microglia, we fit a linear mixed-effects models to test for differences in homeostatic gene expression across experimental conditions, while accounting for technical variation from cell hashing and sample preparation. Statistical comparisons between conditions were performed using estimated marginal means with Benjamini-Hochberg correction for multiple testing.

#### Integration with AD genome-wide association study data

Genetic enrichment analysis was performed using the scDRS v.1.0.2 ^55^ and MAGMA v.1.07a tools ^54^. Initially, gene-level *P* values for the AD GWAS comprising >111,000 clinically diagnosed/proxy AD cases and >677,000 controls ^4^ were computed using the MAGMA *bfile* function. Using the scDRS *munge-gs* function, the top 1,000 AD-associated genes from the MAGMA analysis were extracted, and the gene-level statistics were formatted into an scDRS .gs file. To score each cell on the expression of the top 1,000 AD-associated genes, the scDRS *compute-score* function was run on the raw count matrix with the parameters h5ad-files set to mouse and gs-species set to human. To assess the overall association with AD GWAS signal for each cell population, the scDRS *perform-downstream* function was used. The identified *P* values of associated were corrected for multiple testing using the BH method (adjusting for the number of cell populations).

Subsequently, cell populations with treatment-associated changes in expression of AD-associated genes were identified. For each cell population, a linear mixed-effects model was constructed with the scDRS normalized enrichment score as dependent variable, treatment and time point as independent variables with interaction effects, and mouse and sequencing pool as random effects. This model was constructed separately for veh-PBS and veh-LPS animals and for veh-LPS and sema-LPS animals. Treatment contrasts were tested using a least-squares means two-tailed *t*-test, and *P* values were corrected for multiple testing using the BH method (adjusting for the number of cell populations).

Moreover, for microglia, endothelial cells, and pericytes1, treatment-associated expression levels were identified for the top 50 AD-associated genes. Initially, AD-associated genes were mapped to mouse orthologues. For each cell type and AD-associated gene, a generalized additive model (GAM) was constructed with the raw gene expression as dependent variable, treatment and time point as independent variables with interaction effect, and mouse and sequencing pool as random effects, and *family* set to negative binomial. As above, this model was constructed separately for veh-PBS and veh-LPS animals and for veh-LPS and sema-LPS animals. *P* values were corrected for multiple testing using the BH method (adjusting for the number of tested genes).

#### Integration with AD transcriptional signatures

A list of genes that are up- and downregulated in Class C1 AD individuals (528 and 364 genes, respectively) was obtained from Neff et al. ^60^ and mapped to mouse orthologues. Mouse hippocampal cells were subsequently scored on the activity of these two gene sets using the Seurat function *AddModuleScore.* For each cell population and gene set, a linear mixed-effects model was constructed with the gene set activity as the dependent variable, treatment and time point as independent variables with interaction effects, and mouse and sequencing pool as random effects. This model was constructed separately for veh-PBS and veh-LPS animals and for veh-LPS and sema-LPS animals. Treatment contrasts were tested using a least-squares means two-tailed *t*-test, and *P* values were corrected for multiple testing using the BH method (adjusting for the number of cell populations).

#### Analysis of SEA-AD snRNA-seq dataset

Human immune cells from the middle temporal gyrus and dorsolateral prefrontal cortex were obtained from Gabitto et al. ^61^ and processed individually. Normalization of the counts was performed using the Seurat *SCTransform* function. Subsequently, PCA dimensionality reduction was carried out using the *RunPCA* function, and clustering was completed using the *FindNeighbors* and *FindClusters* functions with the resolution parameter set to 0.1. One cluster overlapped with the cells labeled as lymphocytes, and another cluster contained cells labeled as monocytes.

To identify whether the proportion of lymphocytes was altered in individuals with dementia, a logistic regression model was constructed with dementia status as the dependent variable and lymphocyte proportion for each individual as the independent variable. This model was compared to a null model using a likelihood-ratio test.

To identify cells enriching for markers of mouse neutrophils, the top 500 marker genes for mouse neutrophils in the hippocampal atlas were mapped to human gene symbols. The human immune cells were subsequently scored on the activity of this gene set using the Seurat function *AddModuleScore*.

#### SCENIC analysis

Gene regulatory networks were identified using the python v.0.11.0 implementation of the SCENIC tool ^45^. SCENIC was run separately for each non-neuronal cell type. To test for differentially activated regulons across treatments, a linear mixed-effects model was constructed with the SCENIC regulon activity score as dependent variable, treatment and time point as independent variables with interaction effects, and mouse and sequencing pool as random effects. *P* values were corrected for multiple testing using the BH method (adjusting for the number of regulons). Functional gene set enrichment analysis was carried out for each regulon as described above.

#### Cell-cell communication analysis

To estimate the ligand-receptor interactions between non-neuronal cell types within the snRNA-seq data, the gene expression matrix from non-neuronal cells in mice at 2h and 24h post-PBS or LPS treatment was taken. Next, the R package CellChat (v1.6.1) ^49^ was used with the CellChatDB mouse database for the analysis using default values. Additionally, the CellChat function *identifyOverExpressedGenes* was applied to conduct a differential expression analysis of ligand-receptor pairs between the veh-LPS and veh-PBS treatment group as well as the comparison between the sema-LPS and the veh-LPS group. Only ligand-receptor pairs with the same directionality in the regulation were taken. The percentage of cells value was set to 0.25, the natural log fold-change threshold was set to 0.1 with an adjusted *P* value threshold set to 0.05 for both ligand and receptor.

#### Comparing treatment response between the hippocampus and DVC

To correlate the treatment response to LPS and semaglutide between the hippocampus and DVC, the top 100 differentially expressed genes were identified for each brain region and each non-neuronal cell type comparing cells from veh-LPS and veh-PBS mice and from sema-LPS and veh-LPS mice, respectively, across time points. For each differential gene expression analysis, a pseudo-bulk gene expressed matrix was generated by summing the transcript counts for all cells with the same cell type and mouse combination. DESeq2 was then applied on the pseudo-bulk gene expression matrix with treatment group and time point post-PBS or LPS treatment as variables in the design matrix. The top 100 differentially expressed genes were selected based on the BH-adjusted *P* values. To compare the treatment response to either LPS or semaglutide between the hippocampus and DVC, for each shared non-neuronal cell type, the Spearman’s rho of the log fold-changes of the top 100 LPS- or semaglutide-induced genes in each brain region was computed. To compare the LPS- and semaglutide-induced treatment response, for each non-neuronal cell type, the Spearman’s rho of the log fold-changes of the top 100 LPS-induced and the top 100 semaglutide-induced genes was computed.

### *In vivo* data statistical analysis

Statistical analyses for testing changes in the abundance of markers of peripheral inflammation or neuroinflammation were performed by constructing a linear model which was used as input for a least-squares means two-tailed *t*-test (described in detail below).

For assessing the change in relative IBA1 or GFAP abundance at different doses of LPS, a linear model was constructed for each protein (IBA1 or GFAP) and each brain region (hippocampus or entire brain) using the protein percentage as dependent variable and treatment (veh-PBS, veh-LPS-0.05, veh-LPS-0.1, veh-LPS-0.5, and veh-LPS-1.0) and time (2d and 11d after the last LPS injection) as independent variables with interactions effects. Least-squares means two-tailed *t*-test *P* values were corrected for multiple testing using the BH method (adjusting for the numbers of treatment comparisons and time points). Likewise, for assessing the change in relative IBA1 or GFAP in the hippocampus or entire brain following semaglutide treatment, a linear model was constructed for each protein and each brain region using the protein percentage as dependent variable and treatment (veh-LPS and sema-LPS) and time (2d and 11d after the last LPS injection) as independent variables with interaction effects. Least-squares means two-tailed *t*-test *P* values were corrected for multiple testing using the BH method (adjusting for the number of time points).

For assessing the change in peripheral markers of inflammation, a linear model was constructed for each protein using the protein value (pg/mg) as dependent variable and the treatment group (veh-PBS, veh-LPS, and sema-LPS) and time (2h, 24h, or 5d after the last LPS injection) as independent variables with interaction effects. Least-squares means two-tailed *t*-test *P* values were corrected for multiple testing using the BH method (adjusting for the numbers of treatment comparisons and time points).

## Data and code availability

All single-cell expression data will be made available to the reviewers and upon publication. The source code used to analyze the data and produce the statistical figures is available at https://github.com/perslab/Ludwig-AD-2025.

## Acknowledgements

Novo Nordisk Foundation Center for Basic Metabolic Research is an independent Research Center, based at the University of Copenhagen, Denmark and partially funded by an unconditional donation from the Novo Nordisk Foundation (http://www.cbmr.ku.dk/) (Grant number NNF18CC0034900). THP acknowledges the Lundbeck Foundation (Grant number R190-2014-3904) and the Danish Council for Independent Research (Grant number 8045-00091B). The authors thank Lene Wither Takla (Novo Nordisk A/S, Global Drug Discovery, Denmark) for expert technical assistance with conducting the animal studies, Heidi Solvang Nielsen (Novo Nordisk A/S, Global Drug Discovery, Denmark) for dissection of the hippocampus for RNA-seq and histology, as well as Esther Bloem and Susanne Jørgensen for assistance with protein assays (Novo Nordisk A/S, Global Research Technologies, Denmark). The authors acknowledge Gubra A/S for animal study and IHC support, and the Single-Cell Omics Platform (SCOP) at the Novo Nordisk Foundation Center for Basic Metabolic Research for technical expertise and support.

## Author contributions

Conceptualization of project: MQL, DMR, JPW, LBK, THP. Conceptualization of the LPS-induced neuroinflammation model: SNH, AS, DH, LBK. Rodent studies: SNH, AS, DH, KDD, MM, FW, LBK. Immunohistochemistry: SNH, SB, CP, FW. RNA-sequencing experiments: KLE. Data analysis and interpretation: MQL, MAB, DMR, JM, VD, KLE, AMB, KN, JPW. Manuscript: MQL, MAB, DMR, LBK, THP wrote the first draft of the manuscript. All authors provided comments to and approved the final manuscript. T.H.P. is the guarantor of the manuscript.

## Declaration of generative AI and AI-assisted technologies in the writing process

During the preparation of this work the authors used generative AI tools to improve readability and language of the manuscript. After using these services, the authors reviewed and edited the content as needed and take full responsibility for the content of the published article.

## References

1. Jack Jr., C. R., et al. Revised criteria for diagnosis and staging of Alzheimer’s disease: Alzheimer’s Association Workgroup. Alzheimer’s & Dementia 20, 5143–5169 (2024).

2. Bieger, A. et al. Neuroinflammation Biomarkers in the AT(N) Framework Across the Alzheimer’s Disease Continuum. J Prev Alzheimers Dis 10.14283/jpad.2023.54 (2023) doi:10.14283/jpad.2023.54.

3. Kunkle, B. W. et al. Genetic meta-analysis of diagnosed Alzheimer’s disease identifies new risk loci and implicates Aβ, tau, immunity and lipid processing. Nat Genet 51, 414–430 (2019).

4. Bellenguez, C. et al. New insights into the genetic etiology of Alzheimer’s disease and related dementias. Nat Genet 54, 412–436 (2022).

5. Pugazhenthi, S., Qin, L. & Reddy, P. H. Common neurodegenerative pathways in obesity, diabetes, and Alzheimer’s disease. Biochimica et Biophysica Acta (BBA) - Molecular Basis of Disease 1863, 1037–1045 (2017).

6. Marso, S. P. et al. Semaglutide and Cardiovascular Outcomes in Patients with Type 2 Diabetes. New England Journal of Medicine 375, 1834–1844 (2016).

7. Wilding, J. P. H. et al. Once-Weekly Semaglutide in Adults with Overweight or Obesity. New England Journal of Medicine 384, 989–1002 (2021).

8. Kosiborod, M. N. et al. Semaglutide in Patients with Heart Failure with Preserved Ejection Fraction and Obesity. New England Journal of Medicine 389, 1069–1084 (2023).

9. During, M. J. et al. Glucagon-like peptide-1 receptor is involved in learning and neuroprotection. Nat Med 9, 1173–1179 (2003).

10. Bendotti, G. et al. The anti-inflammatory and immunological properties of GLP-1 Receptor Agonists. Pharmacological Research vol. 182 Preprint at 10.1016/j.phrs.2022.106320 (2022).

11. Nørgaard, C. H. et al. Treatment with glucagon-like peptide-1 receptor agonists and incidence of dementia: Data from pooled double-blind randomized controlled trials and nationwide disease and prescription registers. Alzheimer’s & Dementia: Translational Research & Clinical Interventions 8, e12268 (2022).

12. Wium-Andersen, I. K., Osler, M., Jørgensen, M. B. & Rungby, J. Antidiabetic medication and risk of dementia in patients with type 2 diabetes□: a nested case – control study. (2019).

13. Wang, T., Buse, J. B., Pate, V. & Stürmer, T. 1802-PUB: Liraglutide as a Potential Drug Repurposing Candidate for Alzheimer’s Disease and Related Dementia—Real-World Evidence. Diabetes 72, 1802-PUB (2023).

14. Wang, W. et al. Associations of semaglutide with first-time diagnosis of Alzheimer’s disease in patients with type 2 diabetes: Target trial emulation using nationwide real-world data in the US. Alzheimer’s & Dementia n/a, (2024).

15. Novo Nordisk A/S. Evoke phase 3 trials did not demonstrate a statistically significant reduction in Alzheimer’s disease progression. https://www.novonordisk.com/content/nncorp/global/en/news-and-media/news-and-ir-materials/news-details.html?id=916462 (2025).

16. Yun, S. P. et al. Block of A1 astrocyte conversion by microglia is neuroprotective in models of Parkinson’s disease. Nat Med 24, 931–938 (2018).

17. Brundin, L., Bergkvist, L. & Brundin, P. Fire prevention in the Parkinson’s disease brain. Nat Med 24, 900–902 (2018).

18. Park, J.-S. et al. Blocking microglial activation of reactive astrocytes is neuroprotective in models of Alzheimer’s disease. Acta Neuropathol Commun 9, 78 (2021).

19. Li, Z. et al. Systemic GLP-1R agonist treatment reverses mouse glial and neurovascular cell transcriptomic aging signatures in a genome-wide manner. Commun Biol 1–6 (2021) doi:10.1038/s42003-021-02208-9.

20. Wang, L. et al. Semaglutide attenuates seizure severity and ameliorates cognitive dysfunction by blocking the NLR family pyrin domain containing 3 inflammasome in pentylenetetrazole-kindled mice. Int J Mol Med 48, (2021).

21. Rakipovski, G. et al. The GLP-1 Analogs Liraglutide and Semaglutide Reduce Atherosclerosis in ApoE−/− and LDLr−/− Mice by a Mechanism That Includes Inflammatory Pathways. JACC Basic Transl Sci 3, 844–857 (2018).

22. Coenraad, W., et al. The Cardioprotective Effects of Semaglutide Exceed Those of Dietary Weight Loss in Mice With HFpEF. JACC Basic Transl Sci 0, (2023).

23. Sadek, M. A., Kandil, E. A., El Sayed, N. S., Sayed, H. M. & Rabie, M. A. Semaglutide, a novel glucagon-like peptide-1 agonist, amends experimental autoimmune encephalomyelitis-induced multiple sclerosis in mice: Involvement of the PI3K/Akt/GSK-3β pathway. Int Immunopharmacol 115, 109647 (2023).

24. Chen, L., Xu, H., Zhang, C., He, J. & Wang, Y. Semaglutide alleviates early brain injury following subarachnoid hemorrhage by suppressing ferroptosis and neuroinflammation via SIRT1 pathway. Am J Transl Res 16, 1102–1117 (2024).

25. Zhao, L. et al. Pharmacologically reversible zonation-dependent endothelial cell transcriptomic changes with neurodegenerative disease associations in the aged brain. 1–15 (2020) doi:10.1038/s41467-020-18249-3.

26. Zhang, Q. et al. Blocking C3d+/GFAP+ A1 Astrocyte Conversion with Semaglutide Attenuates Blood-Brain Barrier Disruption in Mice after Ischemic Stroke. Aging Dis 13, 943–959 (2022).

27. Strain, W. D. et al. Effects of Semaglutide on Stroke Subtypes in Type 2 Diabetes: Post Hoc Analysis of the Randomized SUSTAIN 6 and PIONEER 6. Stroke 53, 2749–2757 (2022).

28. Rodbard, H. W. et al. Oral Semaglutide Versus Empagliflozin in Patients With Type 2 Diabetes Uncontrolled on Metformin: The PIONEER 2 Trial. Diabetes Care 42, 2272–2281 (2019).

29. Verma, S. et al. Effects of once-weekly semaglutide 2.4 mg on C-reactive protein in adults with overweight or obesity (STEP 1, 2, and 3): exploratory analyses of three randomised, double-blind, placebo-controlled, phase 3 trials. EClinicalMedicine 55, 101737 (2023).

30. Rode, A. K. O. et al. Induced Human Regulatory T Cells Express the Glucagon-like Peptide-1 Receptor. Cells vol. 11 Preprint at 10.3390/cells11162587 (2022).

31. De Barra, C., et al. Glucagon-like peptide-1 therapy in people with obesity restores natural killer cell metabolism and effector function. Obesity n/a, (2023).

32. Wong, C. K. et al. Central glucagon-like peptide 1 receptor activation inhibits Toll-like receptor agonist-induced inflammation. Cell Metab 36, 130–143.e5 (2024).

33. Jin, H., Li, M., Jeong, E., Castro-Martinez, F. & Zuker, C. S. A body–brain circuit that regulates body inflammatory responses. Nature 630, 695–703 (2024).

34. Langfelder, P. & Horvath, S. WGCNA: an R package for weighted correlation network analysis. BMC Bioinformatics 9, 559 (2008).

35. Duan, L. et al. PDGFRb Cells Rapidly Relay Inflammatory Signal from the Circulatory System to Neurons via Chemokine CCL2. Neuron 100, 183–200.e8 (2018).

36. Posner, D. A., Lee, C. Y. C., Portet, A. & Clatworthy, M. R. Humoral immunity at the brain borders in homeostasis. Curr Opin Immunol 76, 102188 (2022).

37. Yao, Z. et al. A taxonomy of transcriptomic cell types across the isocortex and hippocampal formation. Cell 184, 3222–3241.e26 (2021).

38. Dann, E., Henderson, N. C., Teichmann, S. A., Morgan, M. D. & Marioni, J. C. Differential abundance testing on single-cell data using k-nearest neighbor graphs. Nat Biotechnol 40, 245–253 (2022).

39. Wendeln, A.-C. et al. Innate immune memory in the brain shapes neurological disease hallmarks. Nature 556, 332–338 (2018).

40. Gabery, S., et al. Semaglutide lowers body weight in rodents via distributed neural pathways. JCI Insight 5, (2021).

41. Ludwig, M. Q., et al. A genetic map of the mouse dorsal vagal complex and its role in obesity. Nat Metab 3, (2021).

42. Ilanges, A. et al. Brainstem ADCYAP1+ neurons control multiple aspects of sickness behaviour. Nature 609, 761–771 (2022).

43. Cheng, W. et al. NTS Prlh overcomes orexigenic stimuli and ameliorates dietary and genetic forms of obesity. Nat Commun 12, 5175 (2021).

44. Ly, T. et al. Sequential appetite suppression by oral and visceral feedback to the brainstem. Nature 624, 130–137 (2023).

45. Aibar, S. et al. SCENIC: single-cell regulatory network inference and clustering. Nat Methods 14, 1083–1086 (2017).

46. Van de Sande, B. et al. A scalable SCENIC workflow for single-cell gene regulatory network analysis. Nat Protoc 15, 2247–2276 (2020).

47. Boche, D. & Gordon, M. N. Diversity of transcriptomic microglial phenotypes in aging and Alzheimer’s disease. Alzheimer’s & Dementia 18, 360–376 (2022).

48. Gómez Morillas, A., Besson, V. C. & Lerouet, D. Microglia and Neuroinflammation: What Place for P2RY12? Int J Mol Sci 22, (2021).

49. Jin, S. et al. Inference and analysis of cell-cell communication using CellChat. Nat Commun 12, 1088 (2021).

50. Young, K. A., Biggins, L. & Sharpe, H. J. Protein tyrosine phosphatases in cell adhesion. Biochemical Journal 478, 1061–1083 (2021).

51. Yang, A. C. et al. A human brain vascular atlas reveals diverse mediators of Alzheimer’s risk. Nature 10.1038/s41586-021-04369-3(2022) doi:10.1038/s41586-021-04369-3.

52. Sun, N. et al. Human microglial state dynamics in Alzheimer’s disease progression. Cell 186, 4386–4403.e29 (2023).

53. Sun, N. et al. Single-nucleus multiregion transcriptomic analysis of brain vasculature in Alzheimer’s disease. Nat Neurosci 26, 970–982 (2023).

54. de Leeuw, C. A., Mooij, J. M., Heskes, T. & Posthuma, D. MAGMA: Generalized Gene-Set Analysis of GWAS Data. PLoS Comput Biol 11, e1004219 (2015).

55. Zhang, M. J. et al. Polygenic enrichment distinguishes disease associations of individual cells in single-cell RNA-seq data. Nat Genet 54, 1572–1580 (2022).

56. Boles, J. S., Huarte, O. U. & Tansey, M. G. Microfluidics-free single-cell genomics reveals complex central-peripheral immune crosstalk in the mouse brain during peripheral inflammation. bioRxiv 2023.10.05.561054 (2023) doi:10.1101/2023.10.05.561054.

57. Liu, C.-C., Kanekiyo, T., Xu, H. & Bu, G. Apolipoprotein E and Alzheimer disease: risk, mechanisms and therapy. Nat Rev Neurol 9, 106–118 (2013).

58. García-García, V. A., Alameda, J. P., Page, A. & Casanova, M. L. Role of NF-κB in Ageing and Age-Related Diseases: Lessons from Genetically Modified Mouse Models. Cells vol. 10 Preprint at 10.3390/cells10081906 (2021).

59. Lysosome regulation of microglia in Alzheimer’s disease via TFEB–vacuolar ATPase. Nat Neurosci 27, 13–14 (2024).

60. Neff, R. A. et al. Molecular subtyping of Alzheimer’s disease using RNA sequencing data reveals novel mechanisms and targets. Sci Adv 7, (2021).

61. Gabitto, M. I. et al. Integrated multimodal cell atlas of Alzheimer’s disease. Nat Neurosci 10.1038/s41593-024-01774-5 (2024) doi:10.1038/s41593-024-01774-5.

62. Edison, P. et al. Evaluation of liraglutide in the treatment of Alzheimer’s disease. Alzheimer’s & Dementia 17, e057848 (2021).

63. Gejl, M. et al. In Alzheimer’s Disease, 6-Month Treatment with GLP-1 Analog Prevents Decline of Brain Glucose Metabolism: Randomized, Placebo-Controlled, Double-Blind Clinical Trial. Frontiers in Aging Neuroscience vol. 8 Preprint at https://www.frontiersin.org/articles/10.3389/fnagi.2016.00108 (2016).

64. Frederiksen, K. S. et al. Effects of Semaglutide on Alzheimer’s Disease-Related Biological Processes: Results from a Biofluid Biomarker and Multiomics Immunophenotyping Phase 3 Study in Patients with Early Alzheimer’s Disease After 12 Weeks of Treatment. *https://sciencehub.novonordisk.com/congresses/ctad2025/frederiksen.html* (2025).

65. Tang, B. et al. Genetic Variation in Targets of Antidiabetic Drugs and Alzheimer Disease Risk. Neurology 99, e650 LP-e659 (2022).

66. von Scholten, B. J. et al. Effects of liraglutide on cardiovascular risk biomarkers in patients with type 2 diabetes and albuminuria: A sub-analysis of a randomized, placebo-controlled, double-blind, crossover trial. Diabetes Obes Metab 19, 901–905 (2017).

67. Zobel, E. H. et al. Effect of liraglutide on expression of inflammatory genes in type 2 diabetes. Sci Rep 11, 18522 (2021).

68. Nizari, S. et al. Glucagon-like peptide-1 (GLP-1) receptor activation dilates cerebral arterioles, increases cerebral blood flow, and mediates remote (pre) conditioning neuroprotection against ischaemic stroke. Basic Res Cardiol 1, 1–13 (2021).

69. Ahtiluoto, S. et al. Diabetes, Alzheimer disease, and vascular dementia: A population-based neuropathologic study. Neurology 75, (2010).

70. Butterfield, D. A. & Halliwell, B. Oxidative stress, dysfunctional glucose metabolism and Alzheimer disease. Nature Reviews Neuroscience vol. 20 Preprint at 10.1038/s41583-019-0132-6 (2019).

71. Carranza-Naval, M. J. et al. Liraglutide Reduces Vascular Damage, Neuronal Loss, and Cognitive Impairment in a Mixed Murine Model of Alzheimer’s Disease and Type 2 Diabetes. Front Aging Neurosci 13, (2021).

72. Chen, B. et al. GLP-1 programs the neurovascular landscape. Cell Metab 36, 2173–2189 (2024).

73. Sung, Y. J. et al. Proteomics of brain, CSF, and plasma identifies molecular signatures for distinguishing sporadic and genetic Alzheimer’s disease. Sci Transl Med 15, eabq5923 (2023).

74. Dong, Y. et al. Neutrophil hyperactivation correlates with Alzheimer’s disease progression. Ann Neurol 83, 387–405 (2018).

75. Luo, J., Thomassen, J. Q., Nordestgaard, B. G., Tybjærg-Hansen, A. & Frikke-Schmidt, R. Blood Leukocyte Counts in Alzheimer Disease. JAMA Netw Open 5, e2235648–e2235648 (2022).

76. Illumina. bcl2fastq2 Software v2.19.1. Release notes 1–3 https://emea.support.illumina.com/content/dam/illumina-support/documents/downloads/software/bcl2fastq/bcl2fastq-2-19-1-release-notes-1000000035330-00.pdf (2017).

77. Patro, R., Duggal, G., Love, M. I., Irizarry, R. A. & Kingsford, C. Salmon provides fast and bias-aware quantification of transcript expression. Nat Methods 14, 417–419 (2017).

78. Srivastava, A., Malik, L., Smith, T., Sudbery, I. & Patro, R. Alevin efficiently estimates accurate gene abundances from dscRNA-seq data. Genome Biol 20, 65 (2019).

79. Hao, Y. et al. Integrated analysis of multimodal single-cell data. Cell 184, 3573–3587.e29 (2021).

80. Lun, A. T. L. et al. EmptyDrops: distinguishing cells from empty droplets in droplet-based single-cell RNA sequencing data. Genome Biol 20, 63 (2019).

81. Germain, P.-L., Lun, A., Meixide, C. G., Macnair, W. & Robinson, M. D. Doublet identification in single-cell sequencing data using scDblFinder [version 2; peer review: 2 approved]. F1000Res 10, (2022).

82. Petukhov, V. et al. Case-control analysis of single-cell RNA-seq studies. bioRxiv 2022.03.15.484475 (2022) doi:10.1101/2022.03.15.484475.

