## Supplemental Figures for "Semaglutide Attenuates Neuroinflammation in Mice"

### Supplementary figures

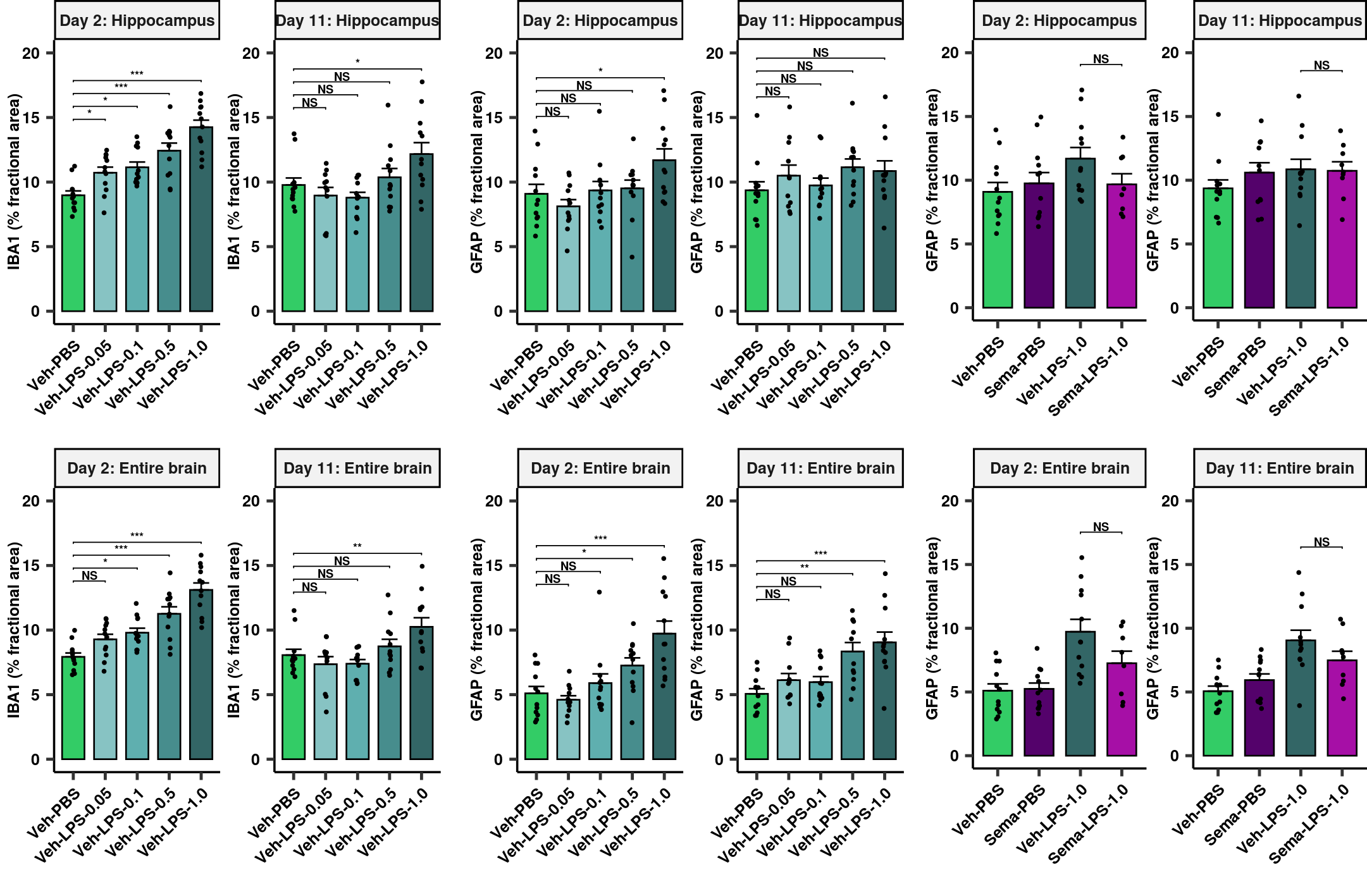

**Supplementary Figure 1. Markers of neuroinflammation in the hippocampus and entire brain follow LPS and semaglutide treatment**

**a**, Dose response to LPS treatment measured as IBA1 quantified by morphometry (fractional area) in the hippocampus and entire brain on termination day 19 and 28 into the study (2 days and 11 days post-LPS administration), respectively. Dots represent individual animals and bars and error bars represent the mean (n=8-12) + SEM. Linear model with BH-adjusted least-squares means two-tailed t-test *P*. **b**, Dose response to LPS treatment measured as GFAP quantified by morphometry (fractional area) in the hippocampus and entire brain on termination day 19 and 28 into the study (2 days and 11 days post-LPS administration), respectively. Dots represent individual animals and bars and error bars represent the mean (n=8-12) + SEM. Linear model with BH-adjusted least-squares means two-tailed t-test P. **c**, Relative GFAP quantified by morphometry (fractional area) in the hippocampus and entire brain on termination day 19 and 28 into the study (2 days and 11 days post-LPS administration), respectively. Dots represent individual animals and bars and error bars represent the mean (n=8-12) + SEM. Linear model with BH-adjusted least-squares means two-tailed t-test *P*. **P*<0.05, ***P*<0.01, ****P*<0.001.

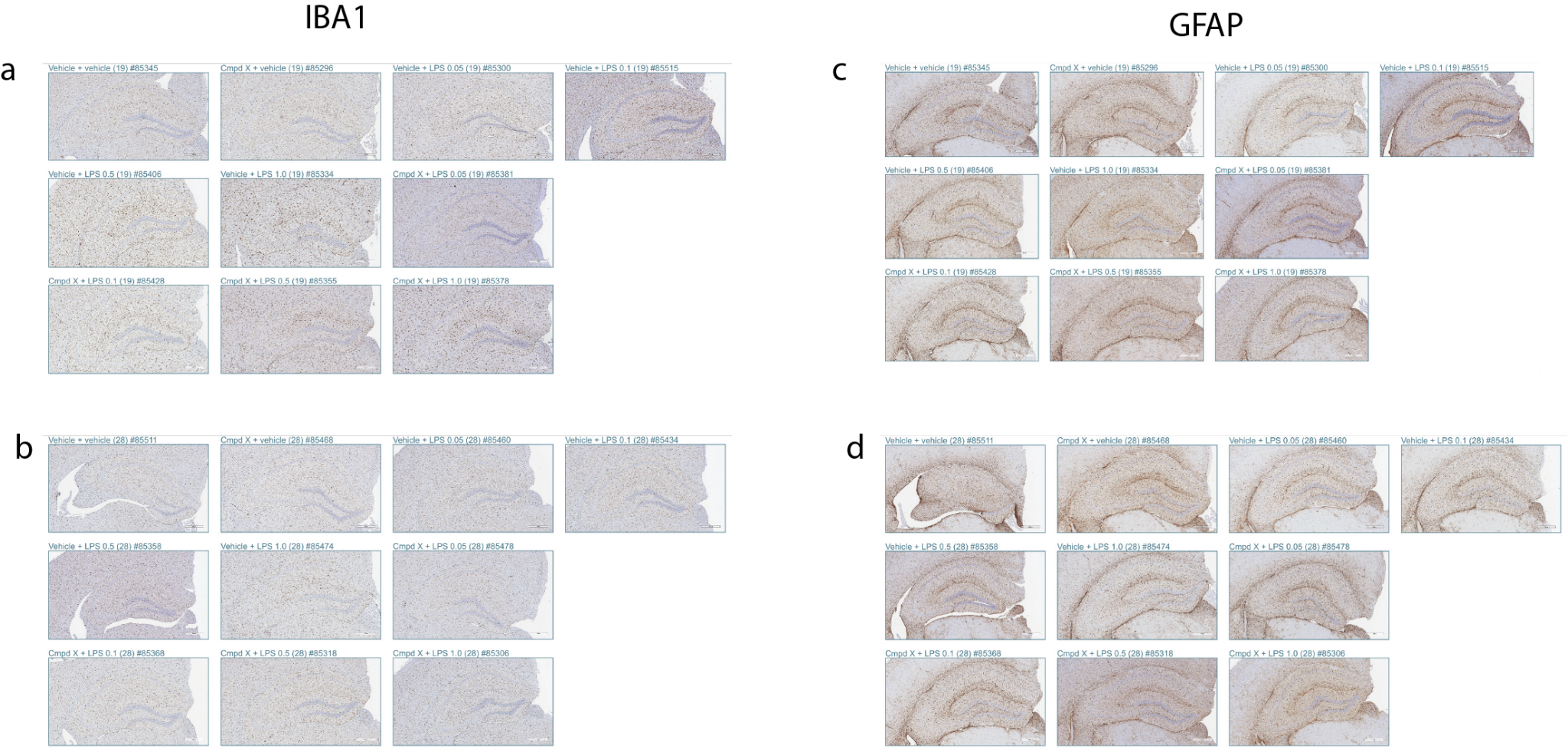

**Supplementary Figure 2. Markers of neuroinflammation in the hippocampus and entire brain follow LPS and semaglutide treatment a**, Representative images of brain sections stained with anti-Iba1 (AbCam, cat. no.ab178847) at day 2 (magnification 5x, scale bar = 400 μm). **b**, Representative images of brain sections stained with anti-Iba1 (AbCam, cat. no.ab178847) at day 11 (magnification 5x, scale bar = 400 μm). **c,** Representative images of brain sections stained with anti-GFAP (Dako, cat. no.Z0334) at day 2 (magnification 5x, scale bar = 400 μm). **d,** Representative images of brain sections stained with anti-GFAP (Dako, cat. no.Z0334) at day 11 (magnification 5x, scale bar = 400 μm). Cmpd X = Semaglutide.

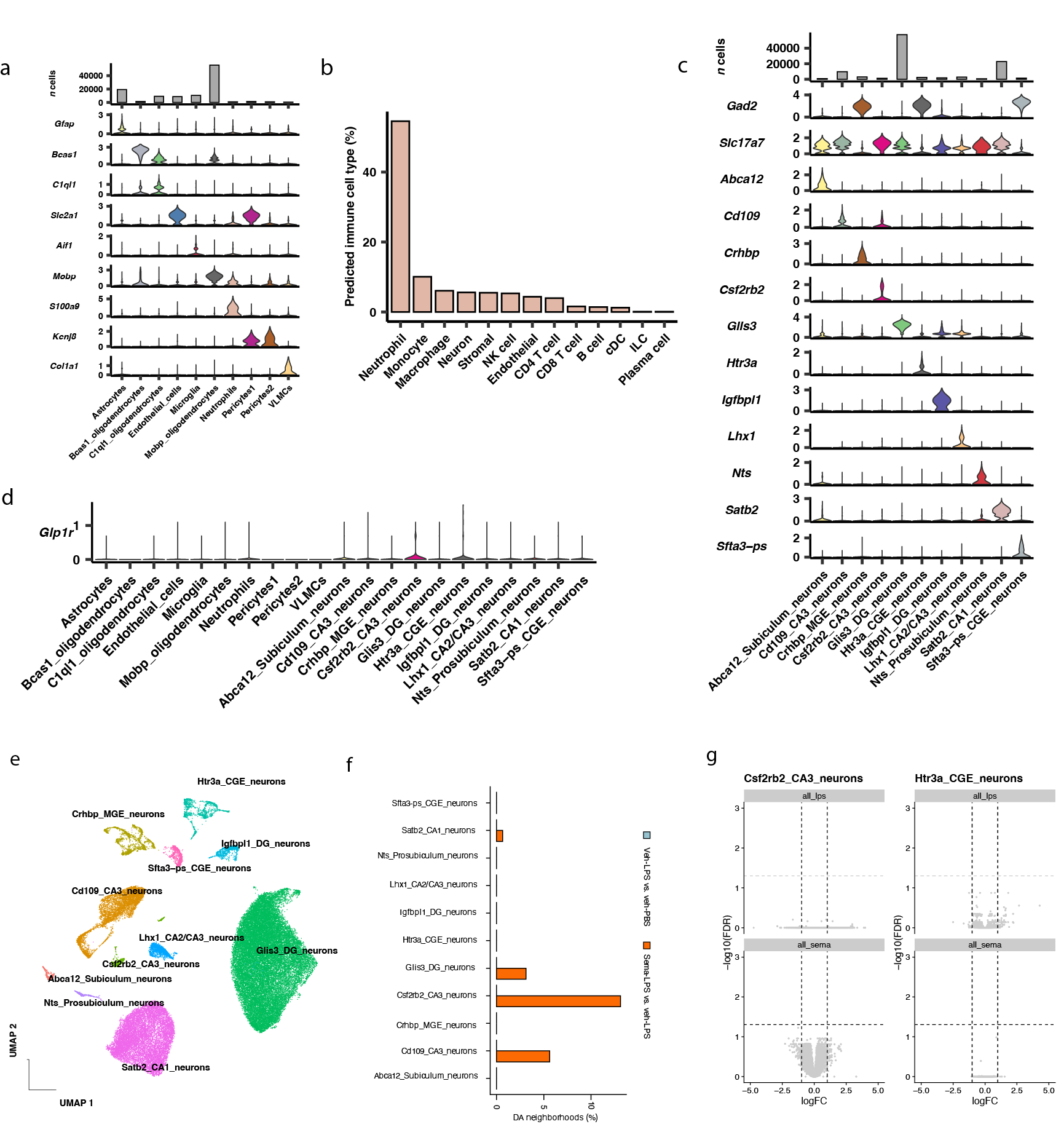

**Supplementary Figure 3. Transcriptomics signatures of hippocampal glial and neuronal populations**

**a**, Stacked barplot indicating number of cells per celltype at top with violin plots of normalized transcript counts for glial marker genes below. **b**, Projection of labels from a consensus sc/snRNA-seq atlas of CNS-border immune cells^1^ to the novel immune cell type identified in the mouse hippocampal atlas. **c**, Stacked barplot indicating number of cells per celltype at top with violin plots of normalized transcript counts for neuronal marker genes. **d,** Violin plot of Glp1r expression across hippocampal neuron cell-types. **e,** UMAP of hippocampal neurons colored by cluster. **f,** , Percentage of cells assigned to differential abundant neighborhoods (changes induced by LPS and semaglutide treatment in turquoise and red, respectively. **f,** Volcano plot of differentially expressed genes in Csf2rb2 (left) and Htr3a (right) neurons in response to LPS (top) or semaglutide (bottom) treatment respectively.

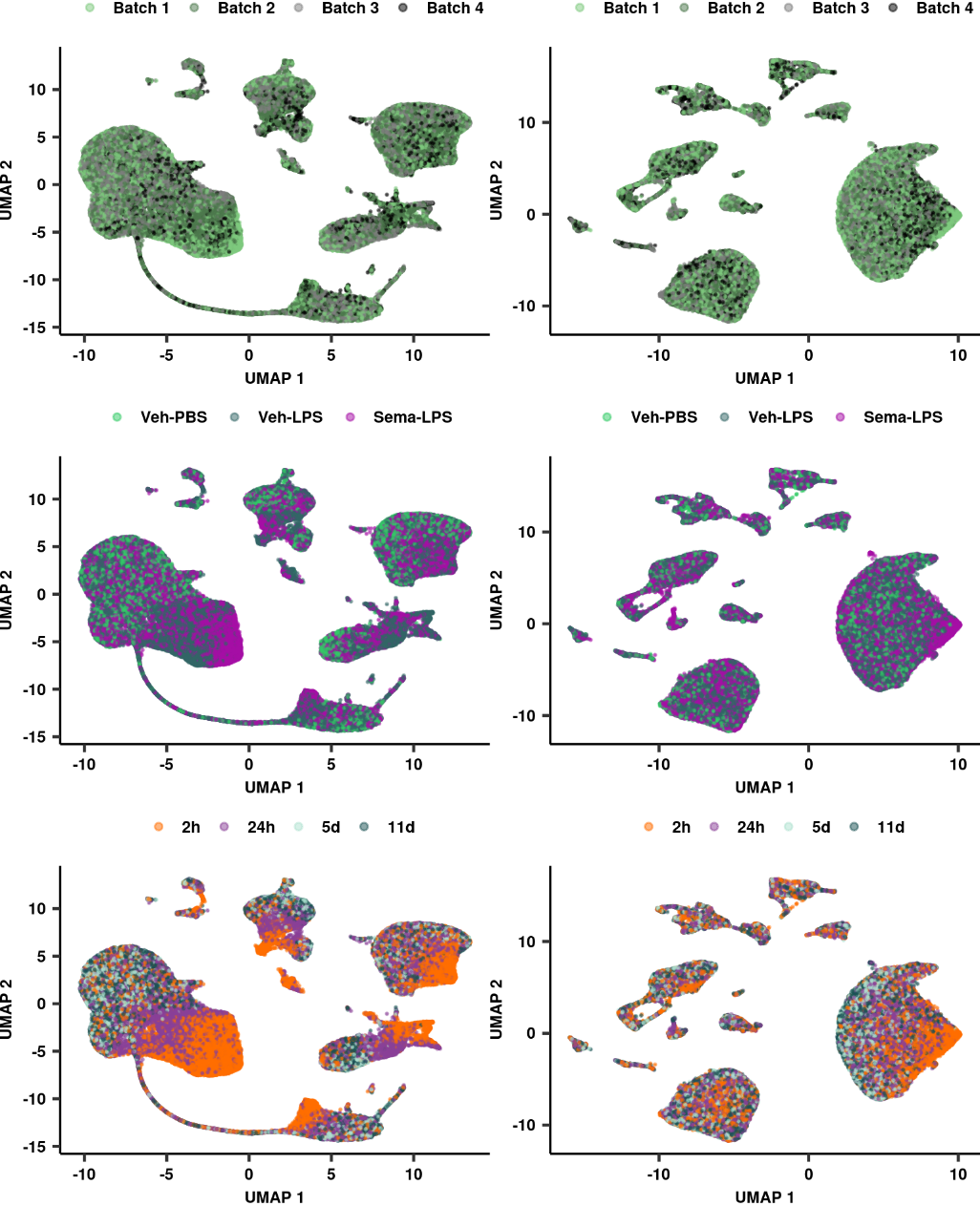

**Supplementary Figure 4. Dimensionality reduction of hippocampal glial cells and neurons**

**a,b,** UMAP of hippocampal glial cells (a) and neurons (b) colored by sequencing batch. **c,d,** UMAP of hippocampal glial cells (c) and neurons (d) colored by treatment group. **e,f,** UMAP of hippocampal glial cells (e) and neurons (f) colored by duration post-LPS or vehicle injection.

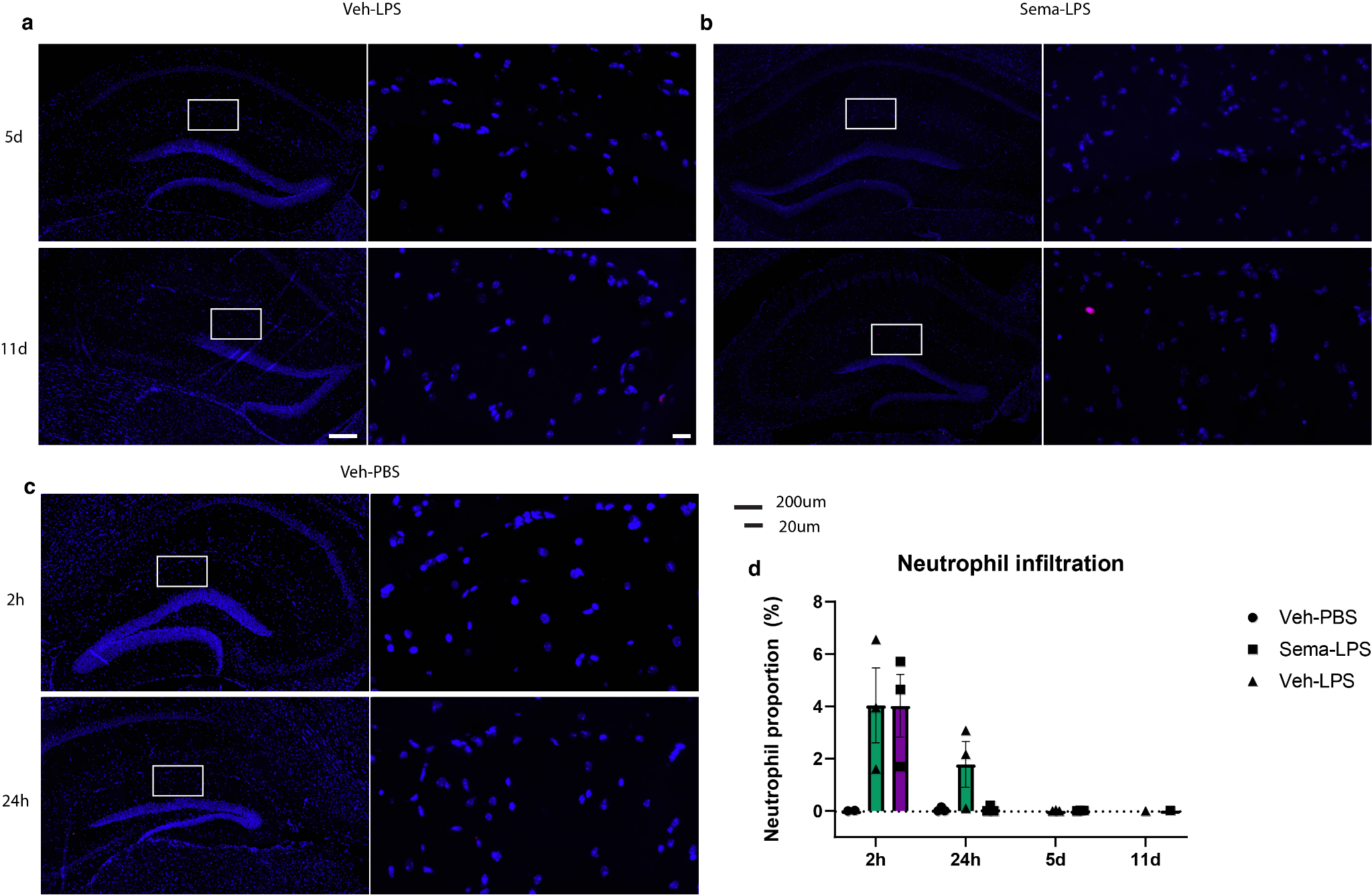

**Supplementary Figure 5. LPS treatment spurs infiltration of neutrophils into the hippocampus**

**a-c,** Immunostaining for anti-neutrophil elastase (NE) (Abcam, Cat. Ab310335) (red) and DAPI (blue). Representative images from Veh-LPS **(a)**, Sema-LPS **(b)**, and Veh-PBS **(c)** at 5 and 11d after final dose of LPS. **e,** Quantification of neutrophil proportion (NE-positive cell count/total cell count) across all time points.

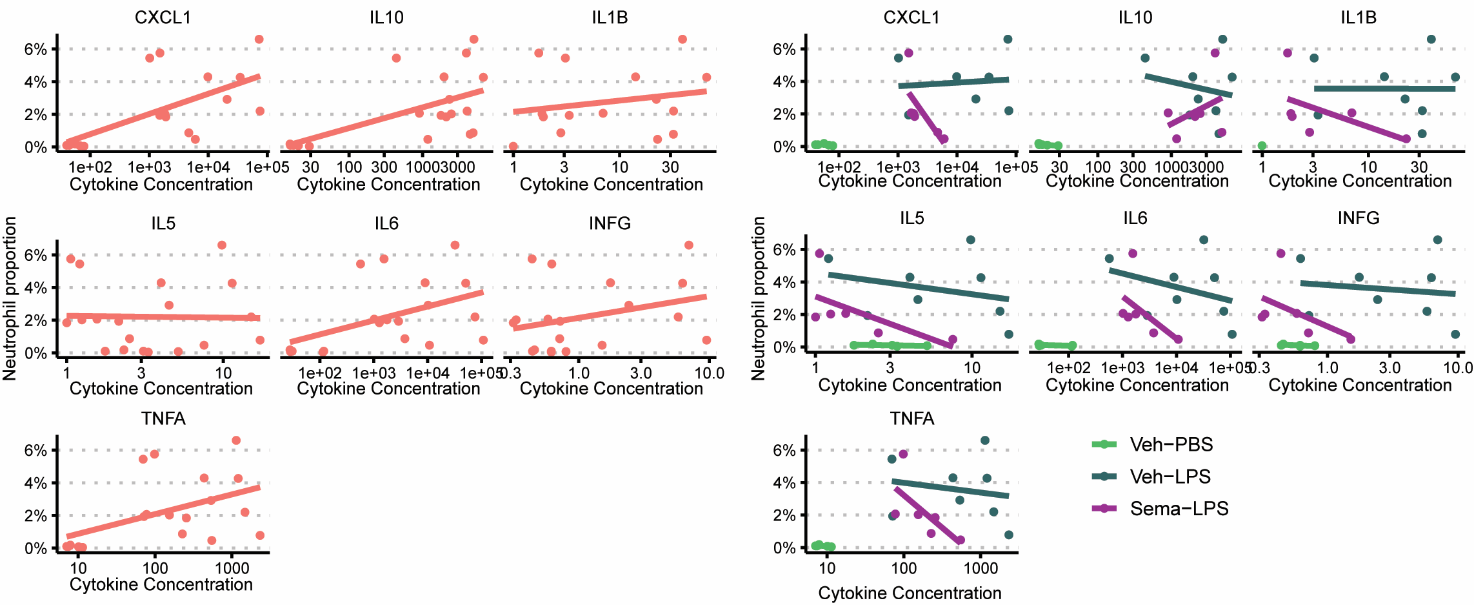

**Supplementary Figure 6. Correlation between cytokine levels and proportional neutrophil infiltration.**

**a,b** Correlation plots of neutrophil proportion (%) and concentration of various cytokines 2h post-LPS for all treatment groups (a) and split according to groups (b). CXCL1, C-X-C motif chemokine ligand 1; INFγ, interferon gamma; IL-1β, interleukin-1 β; IL-5, interleukin-5; IL-6, interleukin-6; IL-10, interleukin-10; TNFα, tumor necrosis factor alpha.

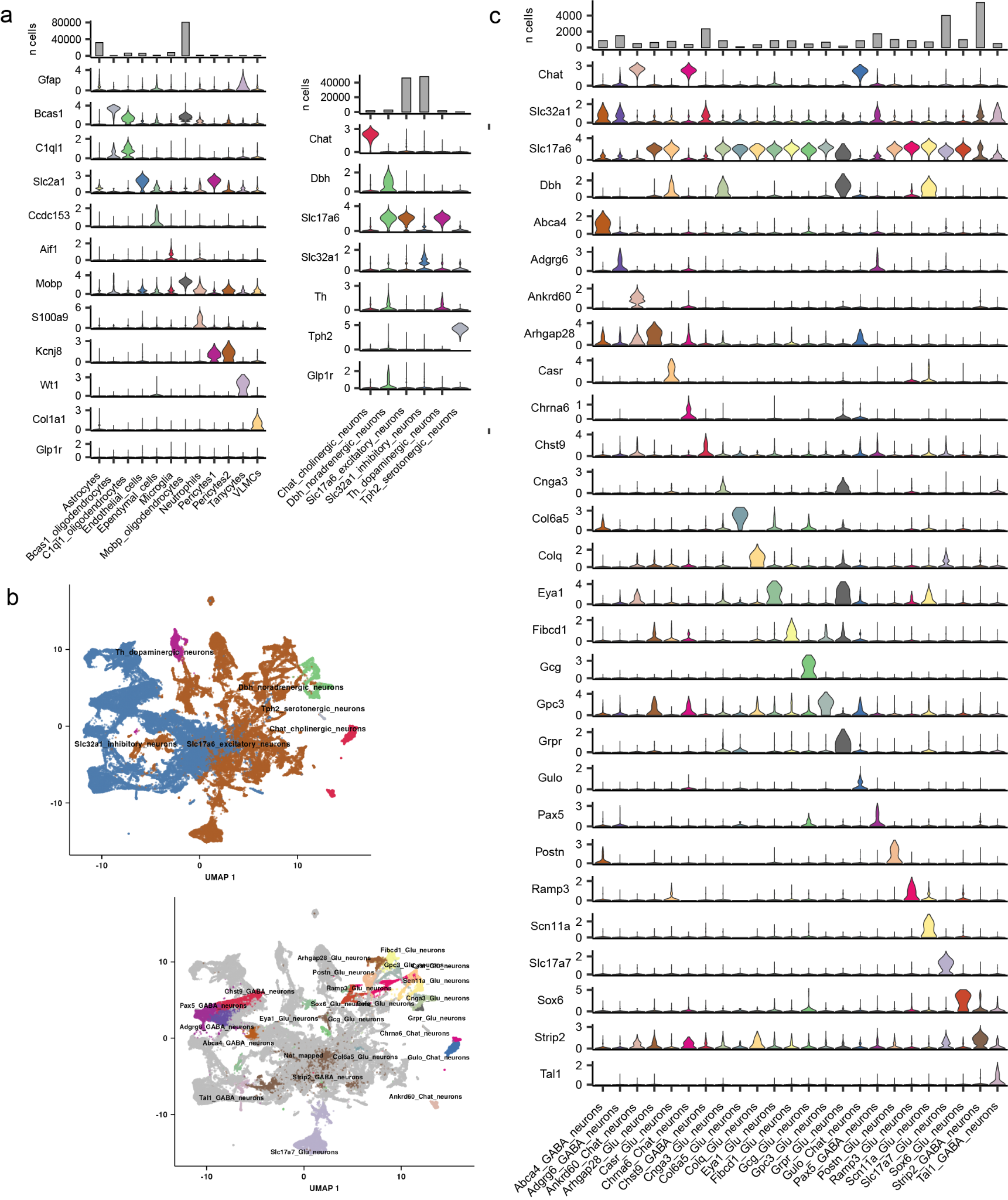

**Supplementary Figure 7. Transcriptomics signatures of DVC glial and neuronal populations**

**a**, Violin plots of normalized transcript counts for glial marker genes (left) and violin plots of normalized transcript counts for neurotransmitter marker genes (right). **b,** UMAP of DVC neurons annotated by high level neurotransmitter clusters (left) and AP centric cluster labels (right). Cells which did not receive an AP centric label are colored in grey. **c**, Violin plot of normalized transcript counts for *Glp1r*.

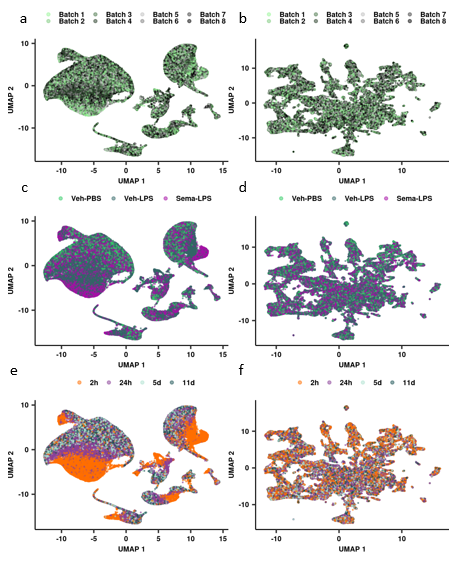

**Supplementary Figure 8. Dimensionality reduction of DVC glial cells and neurons**

**a**,**b**, UMAP of DVC glial cells (a) and neurons (b) colored by sequencing batch. **c**,**d**, UMAP of DVC glial cells (c) and neurons (d) colored by sequencing treatment. **e**,**f**, UMAP of DVC glial cells (e) and neurons (f) colored by duration post-LPS or vehicle injection.

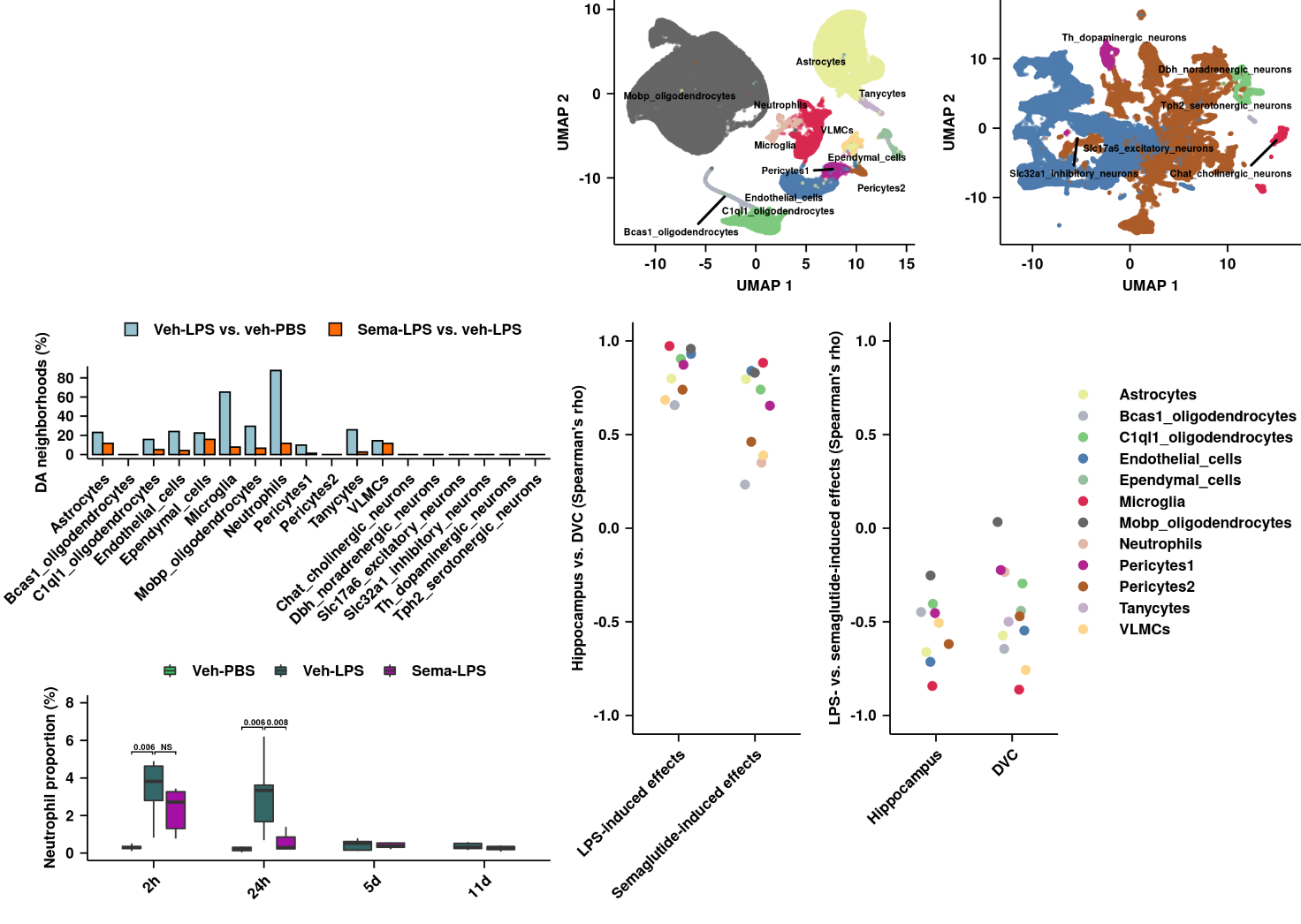

**Supplementary Figure 9. Comparison of the LPS- and semaglutide-induced effects between the hippocampus and DVC**

Spearman’s correlation of the log fold-changes for the top 100 LPS- and the top 100 semaglutide-induced genes between the LPS- and semaglutide-induced effects in hippocampal and DVC non-neuronal cell types.

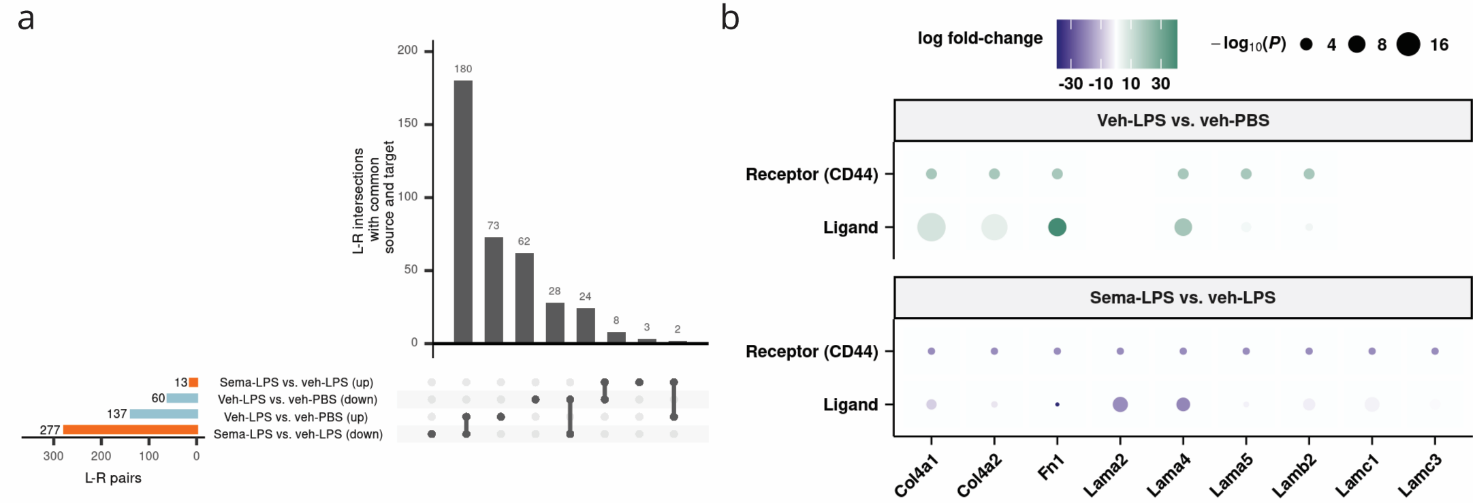

**Supplementary Figure 10. Semaglutide hinders LPS-induced and receptor-ligand interactions**

**a**, Upset plot of the number of differential interactions between a source (expressing the ligand) and a target (expressing the receptor), detected by CellChat comparing veh-LPS vs. veh-PBS and sema-LPS vs. veh-LPS mice. The analysis focuses on microglia, endothelial cells, pericytes1 and astrocytes. LPS, lipopolysaccharide; L-R pairs, ligand-receptor pairs; PBS, phosphate-buffered saline; Sema, semaglutide; Veh, vehicle.

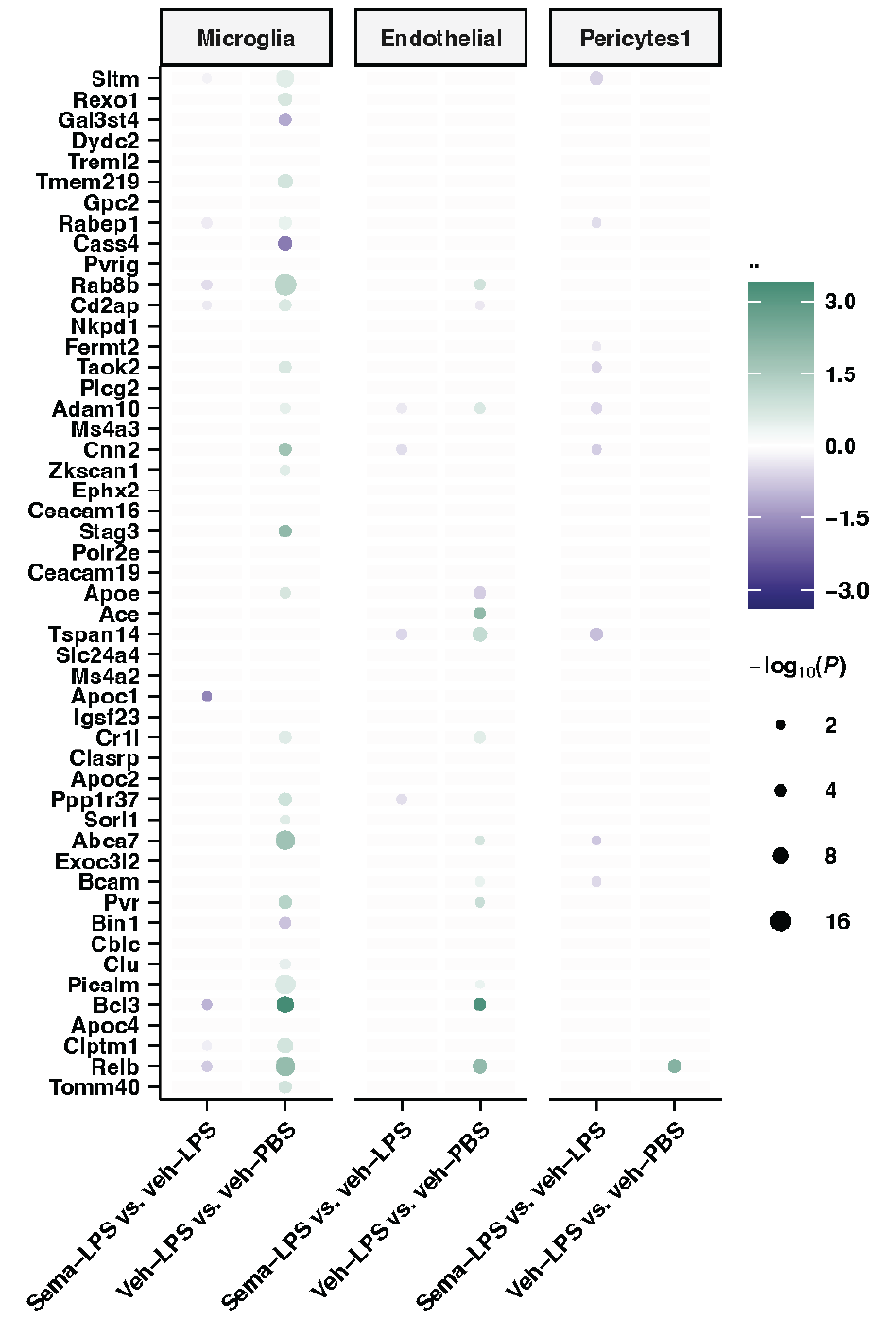

**Supplementary Figure 11. Integration of mouse transcriptomics with AD GWAS data**

**a**, Expression of top 50 AD-associated risk genes in endothelial cells, microglia, and pericytes1 comparing cells from veh-LPS vs. veh-PBS mice and sema-LPS vs. veh-LPS mice. Genes are ranked by their AD GWAS association. Generalized additive model β and BH-adjusted P are shown.

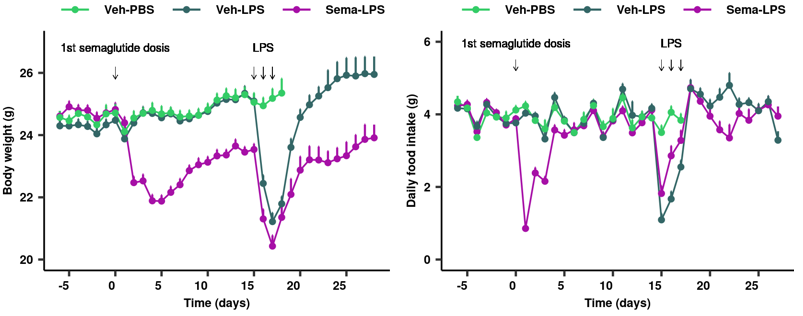

**Supplementary Figure 12. In vivo data for the single-nucleus transcriptomics experiment**

**a**, Body weight following LPS and semaglutide treatment. **b**, Food intake following LPS and semaglutide treatment.

### Supplementary references

1.	Posner, D. A., Lee, C. Y. C., Portet, A. & Clatworthy, M. R. Humoral immunity at the brain borders in homeostasis. *Curr Opin Immunol* **76**, 102188 (2022).

2.	Ludwig, M. Q. *et al.* A genetic map of the mouse dorsal vagal complex and its role in obesity. *Nat Metab* **3**, (2021).

3.	Ilanges, A. *et al.* Brainstem ADCYAP1+ neurons control multiple aspects of sickness behaviour. *Nature* **609**, 761–771 (2022).
